# Synergy between Group 2 capsules and lipopolysaccharide underpins serum resistance in Extra-intestinal Pathogenic *Escherichia coli*

**DOI:** 10.1101/2024.06.07.597965

**Authors:** Naoise McGarry, Domhnall Roe, Stephen Smith

**Affiliations:** Department of Clinical Microbiology, School of Medicine, Trinity College, Dublin

**Keywords:** Escherichia coli, ExPEC, Serum Resistance, Capsule, LPS, Polysaccharides, Antibiotic resistance

## Abstract

*Escherichia coli (E. coli)* is a major cause of urinary tract infections, bacteraemia, and sepsis. CFT073 is a prototypic, urosepsis isolate of sequence type (ST) 73. This laboratory, among others, has shown that strain CFT073 is resistant to serum, with capsule and other extracellular polysaccharides imparting resistance. The interplay of such polysaccharides remains under-explored. This study has shown that CFT073 mutants deficient in lipopolysaccharide (LPS) O-antigen and capsule display exquisite serum sensitivity. Additionally, O-antigen and LPS outer core mutants displayed significantly reduced surface K2 capsule, coupled with increased unbound K2 capsule being detected in the supernatant. The R1 core and O6 antigen are involved in the tethering of K2 capsule to the CFT073 cell surface, highlighting the importance of the R1 core in serum resistance. The dependence of capsule on LPS was shown to be post-transcriptional and related to changes in cell surface charge. Furthermore, immunofluorescence microscopy suggested that the surface pattern of capsule is altered in such LPS core mutants, which display a punctate capsule expression. Finally, targeting LPS biosynthesis using sub-inhibitory concentrations of a WaaG inhibitor resulted in increased serum sensitivity, antibiotic sensitivity, and reduced capsule in CFT073. Interestingly, the dependency of capsule on LPS has been observed previously in several *Klebsiella pneumoniae* isolates and a neonatal meningitic *E. coli* strain, indicating that the synergy between these polysaccharides is not just strain, serotype or species-specific but may be conserved across several pathogenic Gram-negative species. Therefore, using WaaG inhibitor derivatives or phage-derived depolymerases to target LPS is a promising avenue for co-administration with antibiotics to reduce morbidity and mortality by reducing or eliminating surface capsule.

**Impact statement:** Disseminated infections caused by *E. coli* place a large burden on healthcare systems globally, with incidence as well as antimicrobial resistance on the rise (1, 2). The key to designing successful therapeutic strategies is to understand how and why bacteria can cause infection, with an increased need for alternative/ preventative therapies in the age of antimicrobial resistance. ST131 is the predominant and most significant clonal group of bloodstream isolates currently (3-6). The second-most frequently isolated clonal group in *E. coli* bloodstream infections is that of ST73, into which the prototype used in this study, CFT073, is categorised (3, 5-7). Whilst antimicrobial resistance is not currently widespread in the ST73 clonal group, patterns of drug resistance are emerging (7). The clonal group is also associated with higher virulence scores than ST131, highlighting a need to better understand ST73 virulence to identify potential therapeutic intervention strategies (6). This study has identified a synergy between common virulence factors of sepsis-associated *E. coli*, capsule and LPS, which can be exploited to reduce virulence, a promising prospect in the age of antimicrobial resistance.

**Data summary:** Oligonucleotide primers for mutagenesis as well as schematic representations of gene clusters were designed based on the assembled genome sequence of strain CFT073 (GenBank accession number: AE014075.1).

**The authors confirm all other supporting data, code and protocols have been provided within the article or through supplementary data files.**

## Introduction

Extra-intestinal pathogenic *E. coli* (ExPEC) is a collective term for pathogenic strains of *E. coli* which are responsible for extra-intestinal disease, including meningitis, UTIs and bloodstream infection (8). These strains possess genes encoding specialised virulence factors which enable the bacteria to colonise the extra-intestinal sites, in addition to evasion of the host immune responses (9). ExPEC are distinct from intestinal pathogenic *E. coli* in terms of the virulence factors and mechanisms of pathogenesis predominant amongst strains of these extra-intestinal pathotypes, this can be attributed to the ability of *E. coli* to adapt to its niche (10).

Mammalian serum displays a wide variety of highly effective innate immune responses to ExPEC colonisation and bloodstream infection, including the complement system and lysozyme (9). Thus, in order for ExPEC to successfully colonise the host and survive in the blood, these bacteria encode several virulence factors which confer serum resistance. Extracellular polysaccharide factors such as colanic acid, the LPS component O antigen and the surface-associated polysaccharide capsule (K capsule) comprise the extracellular glycome and have been implicated in the resistance of ExPEC to human serum (9, 11-13). A review previously published by this group explores the bactericidal components of human serum, and how ExPEC overcome these defences, in detail (9).

LPS forms a core part of the Gram-negative cell envelope and is essential for structural integrity as well as virulence, attachment, and adhesion to surfaces (4). LPS is comprised of three compartments, the highly conserved lipid A, oligosaccharide core and the highly immunogenic O antigen (14). The LPS O antigen, in juxtaposition to the conserved lipid A, is varied and is used as a serotyping method for *E. coli*. The O antigen is a highly variable region, differing amongst strains and thus, can contribute to pathogenicity. More than 180 different *E. coli* O serotypes have been identified and characterised, with some O antigen types such as O1, O2, O4, O6, O7, O8, O16, O18, O25, and O75 commonly identified in ExPEC and specifically, UTI-associated isolates (12). The O2, O4, O6, O7 and O25 antigens have been specifically implicated in serum resistance of clinical urosepsis isolates and prototypic strains (4, 12, 15).

In contrast to the variable O antigen, there exists only 5 core types in *E. coli:* K-12, R1, R2, R3 and R4. These core groups are differentiated by their outer core structures (16, 17). The R1-type core is the most frequent amongst *E. coli* strains, present in approximately 68% of sequenced strains (16). Interestingly, different core types are correlated with distinct pathotypes, as is also seen with certain O types (i.e. O25 association with extra-intestinal infections) (18). Shiga-toxin producing *E. coli* isolates are predominantly of the R3 core type, whilst many non-pathogenic, commensal isolates are of the K-12 core type (16). Moreover, the R1 core type strains comprise 100% of phylogenetic group B2 *E. coli*, the phylogroup most associated with extra-intestinal infections, indicating a link between core type and pathogenesis/ site of infection (19).

The exopolysaccharide capsule is a cell surface structure composed of long-chain polysaccharides, encasing many isolates and pathotypes of *E. coli*. This exopolysaccharide is similar to those expressed by *Neisseria meningitidis* and *Haemophilus influenzae* among other virulent pathogens, and in those species, capsule is often associated with disseminated infection, immune evasion, and resistance to phagocytosis (20-22). Due to variations in the capsule composition and structures, more than 80 serologically unique K antigens exist across all strains of *E. coli* (23). The *E. coli* capsule fulfils a variety of roles in addition to acting as a steric barrier to the complement system and shielding against other host defences as mentioned above (9). Additionally, the capsule protects against physical environmental stresses, such as dehydration or desiccation (24). Moreover, secreted *E. coli* polysaccharide capsules can contribute to and/or inhibit biofilm formation, depending on the specific K serotype (25, 26).

Despite the hypervariability, there are only two assembly pathways for *E. coli* capsular polysaccharides, designated as Group 1 and 2 prototypes. Groups 3 and 4 are genetic variants of the Group 2 and 1 systems, respectively (27). Each capsule group encodes a distinctive set of cytosolic and inner-membrane enzymes, which generate a distinct capsule sugar structure, which defines the given K serotype (23). Following biosynthesis, a multiprotein complex translocates the nascent capsule polysaccharide across the inner membrane to the outer envelope, where the capsule structure is assembled on the cell surface (13). Most ExPEC express Group 2 capsules on their surfaces with K antigens K1, K2, K5, K100 and K92 being most prevalent amongst ExPEC isolates (26). Furthermore, the K1, K2, K5 and K92 capsules are not just associated with virulent ExPEC isolates but have been shown to be specifically and significantly implicated in the resistance of the isolates to serum (12, 13, 28, 29).

Whilst components of the capsule transport and biosynthesis operons (such as KpsC, KpsS, KpsT) are outer membrane-associated, the mature K capsular polysaccharide is intimately associated with the outer envelope through physiochemical interactions (30). The specific mechanisms which retain capsular polysaccharides to the cell surface are undefined. However, it is strongly evidenced that K1 and K2 capsules (both Group 2 capsules) interact with surface factors such as Braun’s lipoprotein (K2) and LPS core oligosaccharide (K1) to remain associated with the cell, with mutations in these key envelope components resulting in reduced capsule expression at the cell surface (31-33). Interestingly, interactions between the outer core moiety of LPS and capsule polysaccharides have been observed previously in several *Klebsiella* strains of different serotypes (34, 35), in addition to *E. coli* K1 and K5 (31, 33).

The aim of this study was to determine whether the synergy between LPS and capsule may be conserved and exists beyond *Klebsiellae* and *E. coli* K1 and K5, in addition to constructing a mutant deficient in both polysaccharides to determine the full contribution of the glycome to serum resistance in an ExPEC prototype, CFT073.

## Methods

### Bacterial strains and culture conditions

The bacterial strains and plasmids utilised in this study are listed in Supplementary file S1 and S2. Bacteria were grown overnight (approximately 18 hours) in LB Lennox/ Luria broth (LB) NaCl, 5 g/L, Tryptone, 10 g/L Yeast Extract, 5 g/L or on LB agar (Sigma) at 37°C. Bacteria harbouring temperature sensitive plasmids pCP20 and pKD46 were cultured at 30°C. Broth cultures were shaken at 150 rpm for routine overnight culturing. Where required, antibiotics were added to growth media at the following concentrations: 10 μg/mL, gentamicin; 50 μg/mL, kanamycin; 100 μg/mL, carbenicillin (all Sigma). 4-(2-amino-1,3-thiazol-4-yl) (Fluorochem) was solubilised in DMSO at a stock concentration of 1M and was added to bacterial cultures to the desired final concentration.

### Mutagenesis

Mutants were constructed as per the Lambda (λ)-Red recombination protocol as detailed by Datsenko and Wanner (36). Kanamycin (source; pKD4) and gentamicin (pMH2) resistance genes were amplified through PCR by primers which had been designed to contain homologous flanking sequences to the target genes. Amplicons were purified using the Monarch DNA and PCR Cleanup Kit (New England Biolabs) and precipitated as per the Co-Precipitant Pink (Bioline) protocol before resuspension in 4 μL molecular-grade water (Thermo Fisher). The λ -red recombination genes on the pKD46 vector were induced through the addition of L-arabinose (final concentration 10 mM, Sigma) to CFT073/pKD46 cultures for 1.5 hours at 30°C. The temperature-sensitive pKD46 plasmid was removed from CFT073 through incubation at 37°C. Putative mutants were confirmed through PCR. All mutant strains are listed in S1. Plasmids vectors used to amplify antibiotic resistance cassettes for mutagenesis are listed in Table S2. All oligonucleotides used for recombination and mutant screens are listed in S3. To remove antibiotic resistance cassettes, mutants were transformed with pCP20, which possesses the yeast flippase gene FLP, resulting in a marker-less mutant. The protocol followed for FLP-mediated cassette removal is detailed in (36).

### RNA extraction and RT qPCR

Approximately 1×10^7^ bacterial cells were centrifuged at 12,000 x *g* for 1 minute before beginning RNA extraction as per the Monarch Total RNA Miniprep kit protocol (New England Biolabs). RNA was eluted in 30 μL nuclease-free water (Millipore) and quality and concentration were verified by Qubit and Nanodrop analysis. Nanodrop 260/280 and 260/230 scores were used to ensure RNA purity and lack of contamination by proteins or DNA. Real-time quantitative PCR (RT-qPCR) was used to quantify gene expression in *E coli*. The Luna® Universal One-Step RT-qPCR Kit was used for all RT-qPCR reactions in 20 µL volumes with 10 ng RNA used as template. Standard curves were generated with fix 2-fold serial dilutions of wild-type CFT073 RNA. RT-qPCR reactions were set up in triplicate in MicroAmp Fast Optical 96-well reaction plates (Applied Biosystems) and run on the StepOnePlus Real-Time PCR System (Thermo Fisher). Default ‘Fast’ RT-qPCR and melt curve settings were utilised. All oligonucleotides used for RT-qPCR are listed in S3. Data analysis was carried out using the StepOne software or Prism GraphPad. Housekeeping gene *rplT* was used as an internal control. Relative expression (or ‘RQ’) is calculated automatically by the instrument. RQ is the fold change compared to the calibrator (calibrator is usually the untreated or wild-type sample – the calibrator is indicated in each figure legend). The calibrator has a RQ value of 1 and the values for the other samples are the fold-change relative to the calibrator.

### Co-visualisation of O6 and K2 by SDS-PAGE

To prepare whole-cell lysates for SDS-PAGE, 1mL overnight or exponential cultures of CFT073 strains were centrifugued at 13,000 x *g* and resuspended in either fresh LB, 100% NHS or HIS (Biowest) for 10 minutes at 37°. Cells were then centrifuged at 13,000 x *g* and resuspended in 1X Laemelli (Sigma) to a final concentration of 10 OD_600nm_. Samples were boiled at 100°C for 10 minutes and allowed to cool before adding 20 μg/mL Proteinase K (Sigma) and incubating at 56°C for 1 hour. 20 μL of samples were stored at −20°C or run on a 4-20% TruPAGE Precast gel at 180V in 1X TruPAGE SDS Buffer (both Sigma). Following electrophoresis, gels were stained according to the Pro-Q™ Emerald 300 Lipopolysaccharide Stain Kit (Thermo Fisher) and visualised under the Quantity One® (Bio-Rad) imaging system. Alternatively, gels were stained with 0.125% Alcian blue solution (Sigma) to observe total polysaccharide content.

### Western blot and densitometry

CFT073 whole cell lysates were separated by SDS-PAGE on TruPAGE precast gels as described above. Gels were transferred onto polyvinyl difluoride (PVDF) using the iBlot 2 (Thermo Fisher) for dry transfer. Membranes were blocked in 5% Bovine Serum Albumin (BSA) (Sigma) in 1X PBS supplemented with 0.1% Tween 20 (0.1% PBS-T) for 1 hour at room temperature (RT). Membranes were incubated in anti-O6 or anti-K2 primary antibody (Statens Serum Institut) diluted 1/500 in 5% BSA 0.1% PBS-T for 90 minutes at room temperature (RT). Membranes were washed three times for 10 minutes in 0.1% PBS-T before incubating with anti-rabbit-HRP antibody (Cell Signalling Technologies) diluted 1/10,000 in 5% BSA 0.1% PBS-T for 45 minutes at RT. Membranes were washed 4 times before staining for 5 minutes with Pierce ECL Western Blotting Substrate (Thermo Fisher) as per the manufacturer’s protocol. Membranes were imaged under the ImageQuant Las4000 (GE Healthcare Life Sciences). ImageStudio Lite (v 5.5.4) was utilised to relatively (no absolute values) quantify bands from Western blot .TIFF images. The quantitative values reflect the relative amount of polysaccharide as a ratio of each band relative to the gel background.

### Surface hydrophobicity

Percentage hydrophobicity was determined for CFT073 and derivatives using the protocol described in (37). Briefly, overnight cultures were harvested by centrifugation and washed twice in 0.9% NaCl solution. Cells were standardised to an OD_600nm_ of 1 in 0.9% NaCl. A volume of 300 µl of n-hexadecane was added to 1.4 mL of the above cell suspensions before mixing by vortex. The phases were allowed to separate at room temperature for 30 minutes before the optical density of the aqueous phase was measured at OD_600nm_. Surface hydrophobicity was calculated as the percentage of OD extracted into the n-hexadecane phase.

### ELISA

Cultures were washed twice in PBS and standardised to OD_600_ = 0.1 in PBS before plating 25 μL onto a poly-L lysine treated 96-well plate (Greiner) for overnight incubation at 4°C. The plates were centrifuged for 1 minute at 10,000 x *g* before the removal of supernatant and addition of 100 μL 0.1% v/v glutaraldehyde (Sigma) for 10 minutes at RT. Wells were washed three times with PBS-T before the addition of 200 μL 5% (w/v) skimmed milk (Marvel) in PBS-T). Plates were blocked for an hour at 37°C before the addition of anti-OmpF or anti-K2 antibody diluted 1/1000 to each well for incubation at 37°C for 1 hour. Wells were washed three times vigorously with PBS-T before addition of 200 μL anti-rabbit-AP antibody (Cell Signalling Technologies) at 1/10,000 dilution to each well. Wells were washed for a final three times with PBS-T before adding 100μL of the 1-Step PNPP (Thermo Fisher) to each well. The substrate was mixed thoroughly by gently agitating the plate at RT for 30 minutes. To stop the reaction, 50 μL 2M NaOH was added. Absorbance was measured at 405 nm.

### Immunofluorescence microscopy

To prepare bacterial cultures for immunofluorescence microscopy 1 mL stationary phase or exponential cultures were standardised to 0.1 OD_600nm_ and washed twice in PBS before re-suspending in 100 μL 1X PBS/ 0.01% Glutaraldehyde (Sigma). A loop-full (approx. 5 μL) of culture was spread onto a glass slide before drying at 50°C. 50 μL 5% BSA solution in 0.1% PBS-Tween 20 (Sigma) was added to the slide and incubated at 37°C for 30 minutes before the slide was dried as before. Once dried, 50 μL of anti-K2 or anti-O6 antibody (Statens Serum Institut) was added to the slide, diluted 1/500 in BSA solution before incubating at 37°C for 90 minutes. The antibody solution was washed off the slide three times with PBS-Tween. Slides were dried again at 50°C, prior to the addition of anti-rabbit Alexa-Fluor 594-conjugated secondary antibody (ThermoFisher) diluted 1/10,000 in 5% BSA and incubated at 37°C for 30 minutes. The slide was rinsed three more times before a final drying step. A loop-full of 80% sterile glycerol (v/v) was added to the slide before the addition of the cover slip prior to visualisation and imaging under the EVOS M5000 (ThermoFisher).

### Serum killing assay

The serum sensitivity of CFT073 and mutant derivatives was examined by exposing cells to 50% normal human serum (NHS) (Biowest) and heat-inactivated serum (HIS) for 90 minutes at 37°C and determining survival through viable counting as previously described (11). Serum was heat-inactivated by heating at 56°C for 30 minutes. Percentage survival was calculated as a fraction of CFU/mL at T90 in NHS over CFU/mL at T90 in HIS (x100). Serum was heat inactivated at 56°C for 30 minutes. Serum resistance was measured at logarithmic (OD_600nm_ = 0.6) and stationary phase (OD_600nm_ = 3.0). Standardised assays were conducted in order to directly compare response of stationary phase and logarithmic cells to serum at equivalent cell numbers. Stationary phase cultures (OD_600_ = 3.0) were diluted to OD_600_ = 0.6 in conditioned LB (the cell-free growth supernatant of the strain).

### Minimum inhibitory concentration testing

The agar dilution method involved the addition of varying concentrations of rifampicin (Sigma) to nutrient agar medium (Sigma), prior to polymerisation. The prepared agar plates were then inoculated with 100 μL culture which was spread evenly across the plate using sterile bacteriological swabs, with standardised concentrations of overnight culture obtained through serial dilution following absorbance measurements. Minimum inhibitory concentration (MIC) was determined as the lowest concentration which inhibited bacterial growth.

### Graphing

All graphing was completed using GraphPad Prism (v.9.5.0). Graphical figures were made using Biorender (Biorender, online graphing tool).

### Statistical analysis

All statistical analysis was carried out using the GraphPad Prism software (v.9.5.0) unless otherwise stated. Statistical analysis was performed exclusively on biological replicates, wherein experiments were conducted a minimum of 3 (N = 3) or 4 times (N = 4). In all data, a *p* value of ≤ 0.05 is denoted *; a *p* value of ≤ 0.01 is denoted **; a *p* value of ≤ 0.001 is denoted ***; and a *p* value of ≤ 0.0001 is denoted ****.

## Results

### Mutagenesis and visualisation of the CFT073 extracellular glycome

Components of the extracellular glycome (K2 capsule, LPS outer core and O6 antigen) were mutated with specific genes essential in the biosynthesis of the polysaccharides targeted. The region 2 (serotype specific) operon of the K2 capsule biosynthesis cluster was targeted to make a mutant lacking the K2 capsule, Δ*ksl*, in addition to *kpsC*, a gene essential for capsule export (Figure 1A). Outer core biosynthesis was targeted through deletion of the gene encoding the core glycosylase enzyme WaaG (Figure 1B). Lastly, an O6 antigen-deficient mutant was made through deletion of the O antigen ligase gene, WaaL (Figure 1C). Whole cell lysates of wild-type and the panel of glycome mutants were separated by SDS-PAGE and stained with ProQ Emerald 300 (see Figure 1D). Figure 1D shows the stained extracellular glycome of CFT073, consisting of the full-length, smooth LPS molecule and the high molecular weight K2 capsule, as well as the mutants lacking in the corresponding bands to their mutation, e.g., the *waaL* mutant possesses capsule and lipid A-core but is deficient in O6 antigen due to the deleted O antigen ligase. Interestingly, *waaG* mutants do not appear to possess K2 capsule as evidenced by the stained SDS-PAGE gel.

**Figure 1.**
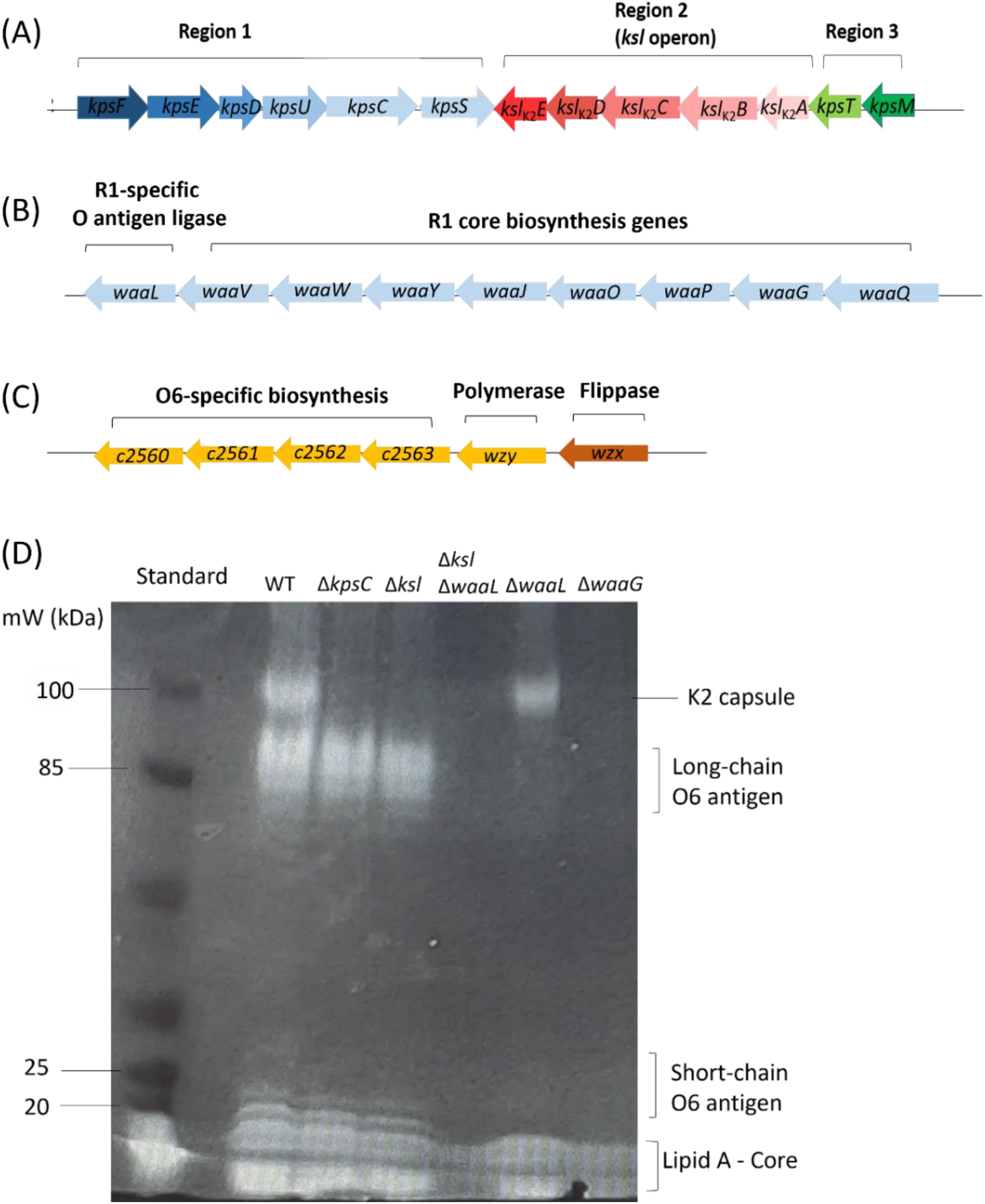
Mutagenesis and visualisation of CFT073 extracellular glycome. (A), (B) and (C) depict the regions targeted in mutagenesis, specifically the K2 capsule gene cluster, the R1 core biosynthesis cluster, and the O6 antigen biosynthesis operon. (D) Whole cell lysates of strain CFT073 and mutant derivatives were separated by SDS-PAGE and stained with the ProQ Emerald 300 LPS stain, allowing co-visualisation of LPS and capsule.

### O antigen and outer core mutants display reduced capsule

Next, a protocol for efficiently blotting the high molecular weight polysaccharides was optimised. Western blot analysis depicted full LPS in strain CFT073 wild-type and *ksl* mutant whole cell lysates, but not in *waaL, waaL/ksl* or *waaG* mutant preparations, as expected (Figure 2A). Similarly, K2 capsule was detected in Western blot analysis of wild-type, *waaL* and *waaG* whole cell preparations, but was not detected in the *kslABCDE* operon mutant, as seen in Figure 2B. Interestingly, there appears to be less visible capsule in O antigen and *waaG* mutants, however, these data are qualitative and require quantification. The same cell preparations used in Figure 2A and B were stained with 0.125% Alcian blue (Figure 2C) following separation by SDS-PAGE to serve as a loading control for the standardised whole cell lysates, showing equivalent polysaccharide in each lane, only with less in the *ksl* mutant lane at around 120 kDa, where the K2 capsule resolves.

**Figure 2.**
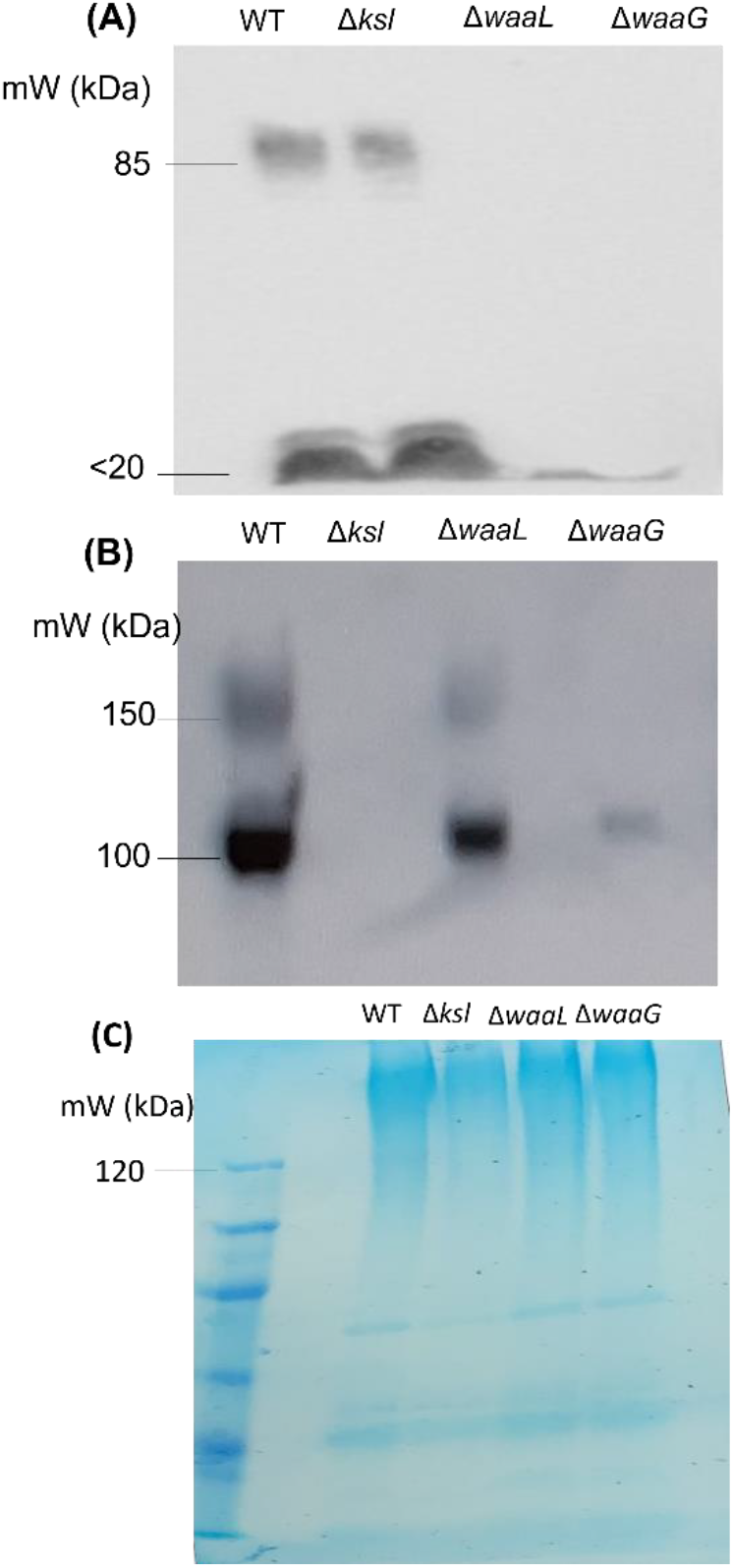
Western blot analyses depict reduced capsule in LPS mutants. Whole cell lysates of strain CFT073 and derivative glycome mutants were separated by SDS-PAGE and subject to (A) Western blot analysis of the O6 antigen, using an O6-specific antibody, (B) Western blot analysis of the K2 capsule, using a K2-specific antibody and (C) a 0.125% Alcian blue total polysaccharide stain, which served as a loading control.

### LPS mutants possess quantifiable reductions to K2 capsule

Next, the apparent reduction in K2 capsule levels observed in LPS mutants was quantified by densitometry (Figure 3A) and ELISA (Figure 3B). Densitometry was performed on N=4 biological replicates of the Western blot performed on whole cell lysates depicted in Figure 2B. The densitometric quantification of K2 capsule showed that *waaL* mutants possess approximately 40% less K2 capsule than wild-type, as detected by Western blot, a statistically significant reduction. Similarly, *waaG* mutants displayed 86% less K2 capsule than wild-type. Interestingly, ELISA quantification of K2 capsule showed significantly reduced capsule in *waaG* mutants compared to wild-type (*p* ≤ 0.05), whilst *waaL* mutants displayed no significant reduction in capsule levels compared to wild-type (*p* = 0.8). The reduction in capsule seen for *waaG* (and *waaL*) mutants was less pronounced when examined by ELISA in comparison to Western blot.

**Figure 3.**
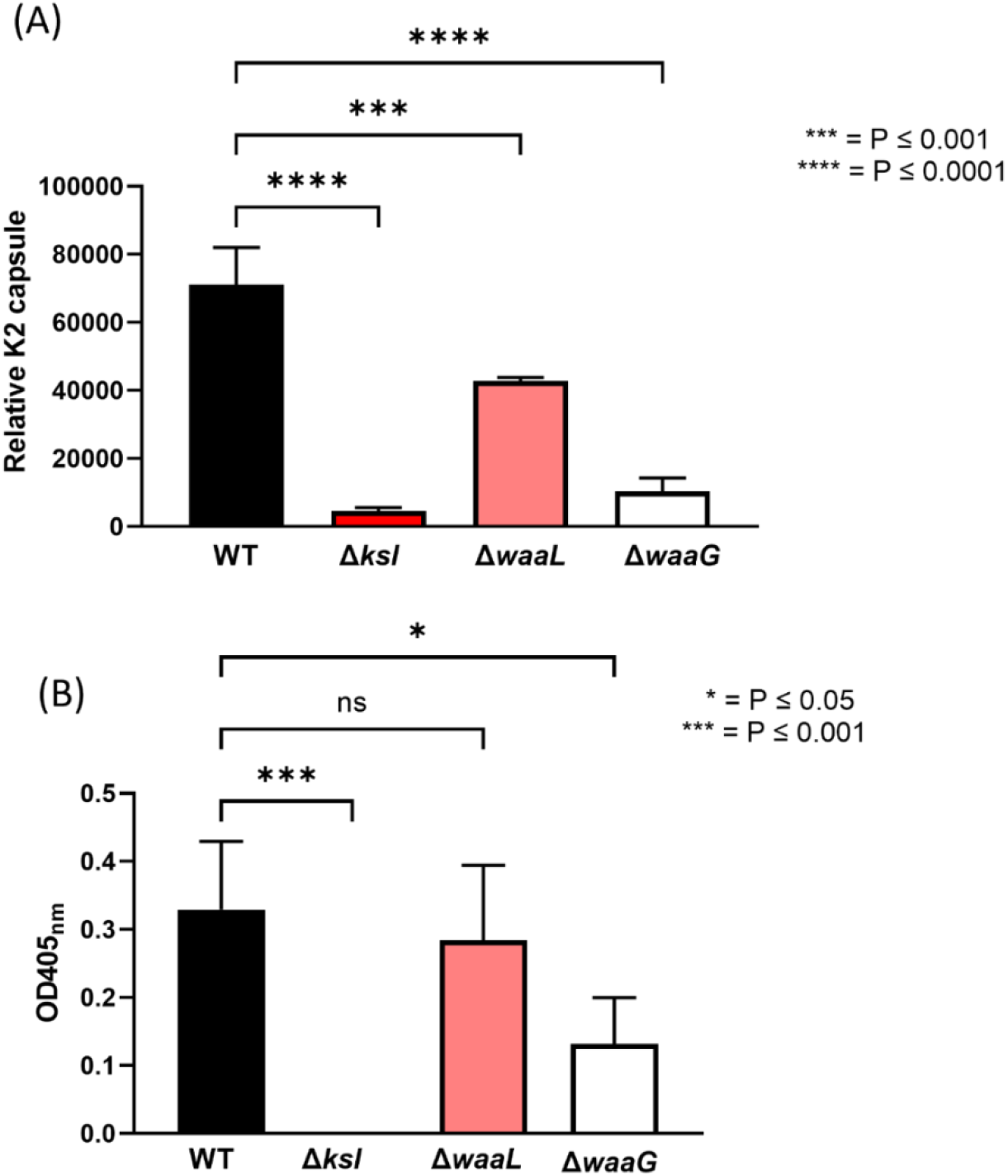
LPS mutants display quantifiable reductions in K2. (A) Densitometric analysis of Western blots performed on whole cell lysates of CFT073 and mutants and (B) ELISA performed on wild-type and mutant cultures allowed for quantification of K2 capsule. Relative K2 capsule: densitometry does not give rise to absolute values. N = 4 for both assays. WT = Wild-type CFT073. Statistical significance calculated by One-way ANOVA and Dunnett’s multiple comparisons.

### LPS mutants have increased capsule in culture supernatants

Due to the unexpected findings depicted in Figure 3B, whereby ELISA quantification showed that LPS mutants did not display pronounced reductions in capsule, it was hypothesised that the anti-K2 antibodies were binding secreted K2 capsule. The ELISA protocol used in this study includes the incubation of cells prior to fixing, allowing for the secretion of molecules, whilst whole-cell lysate preparation for Western blot involves the removal of culture supernatant prior to SDS-PAGE. Culture supernatants taken from strain CFT073 and mutant derivatives were concentrated 10-fold prior to SDS-PAGE and Western blot analysis. As depicted in Figure 4A, the anti-K2 Western blot confirmed K2 capsule in the supernatant of wild-type, *waaL* and *waaG* mutants. Next, the secreted K2 in culture supernatants was quantified by ELISA, as shown in Figure 4B. The *waaL* mutant displayed more secreted capsule in culture supernatant than wild-type, although this increase was not statistically significant (*p* = 0.4), whilst the *waaG* mutant displayed significantly more secreted K2 than wild-type (*p* = 0.0001). These results indicate that the reduced capsule seen in LPS mutants occurs due to changes in surface charge, which has an adverse effect on capsule surface retention.

**Figure 4.**
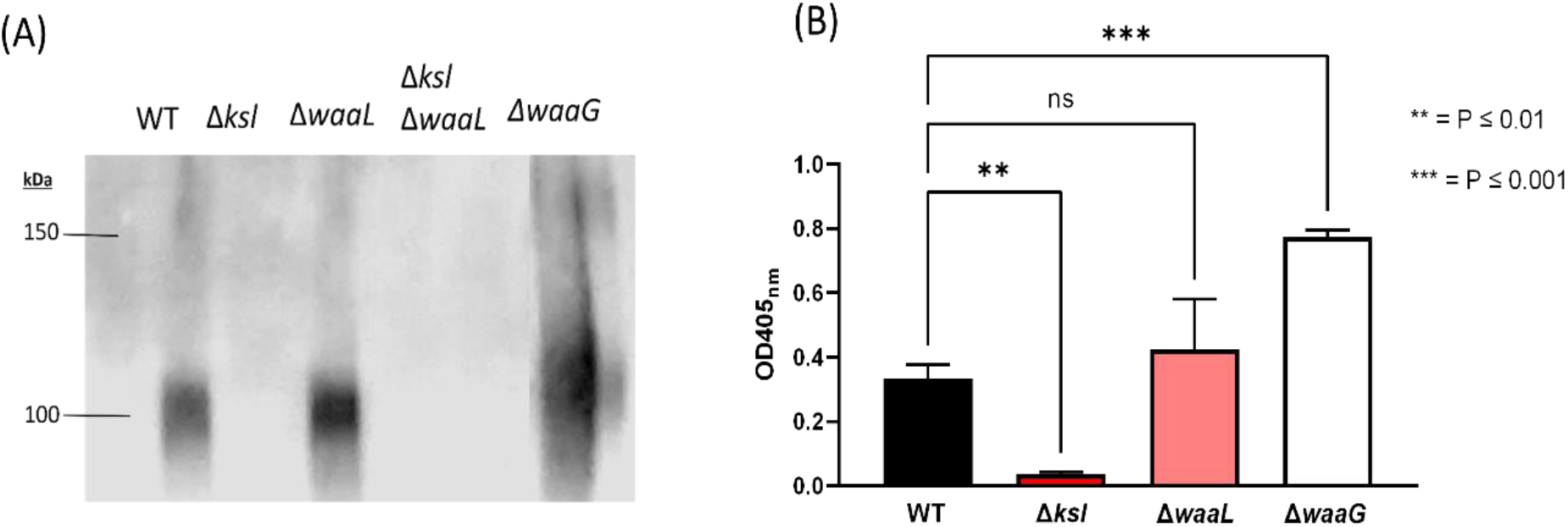
LPS mutants possess increased K2 in the supernatant. (A) Western blot analysis of K2 capsule in stationary phase supernatant taken from bacterial cultures and concentrated 10-fold. (B) ELISA quantification of K2 capsule in stationary phase supernatant of CFT073 and mutants. N = 4 for both assays. WT = Wild-type CFT073. Statistical significance calculated by One-way ANOVA and Dunnett’s multiple comparisons.

### Reduced capsule seen in LPS mutants is post-transcriptional

Following the observation that LPS mutants display reduced surface capsule and increased secreted capsule, it was hypothesised that the dependence of capsule on full-length LPS was post-transcriptional and related to changes in surface charge. RT-qPCR was conducted on RNA extracted from strain CFT073 wild-type, as well as *ksl* (not shown), *waaL* and *waaG* mutants to compare relative expression of the *kpsT/ksl2A* genes. As shown in Figure 5, *waaL* and *waaG* mutants displayed no changes to *kpsT/ksl2A* gene transcription when compared to wild-type. Overall, these results indicate that there are no downstream effects on K2 gene expression in LPS mutants.

**Figure 5.**
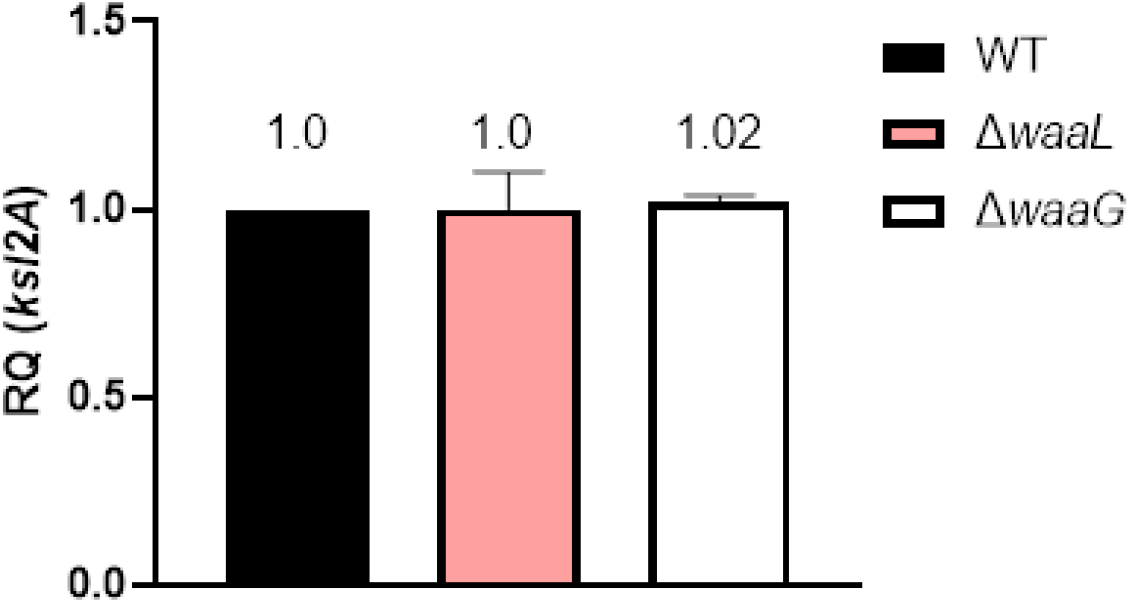
Reduced capsule in LPS mutants occurs post-transcriptionally. RNA was extracted from stationary phase cultures grown under standard lab conditions (LB, 37°C) and *ksl2A* transcript levels were compared. Data analysis and relative expression calculation was carried out automatically on the StepOne software, with housekeeping gene *rplT* used as an internal control. Statistical analysis by One-way ANOVA and Dunnett’s multiple comparisons.WT = Wild-type CFT073. N=4 biological replicates.

### Complementation of LPS restores capsule levels

In order to confirm that the synergy between LPS and capsule is related to surface interactions and is not due to downstream or compensatory mutations, the *waaG* mutant was complemented with pBAD-His-WaaG, a vector with inducible *waaG* expression. Figure 6 shows Western blot analysis of K2 expression in wild-type and Δ*waaG*, in addition to the complemented *waaG* mutant under induced (+ L-arabinose) and un-induced conditions. As seen in Figure 6, the *waaG* mutant complemented with pBAD-His-WaaG post-induction with L-arabinose displayed restored K2 capsule expression, confirming that the *waaG* mutation is non-polar and that the synergy between LPS and capsule occurs due to electrostatic reactions at the cell surface. The O6 antigen mutant deficient in *waaL* also displayed restored K2 levels comparable to that of wild-type upon complementation of the *waaL* gene (data not shown), indicating that both moieties of LPS (O antigen and core) interact with K2 to facilitate surface retention.

**Figure 6.**
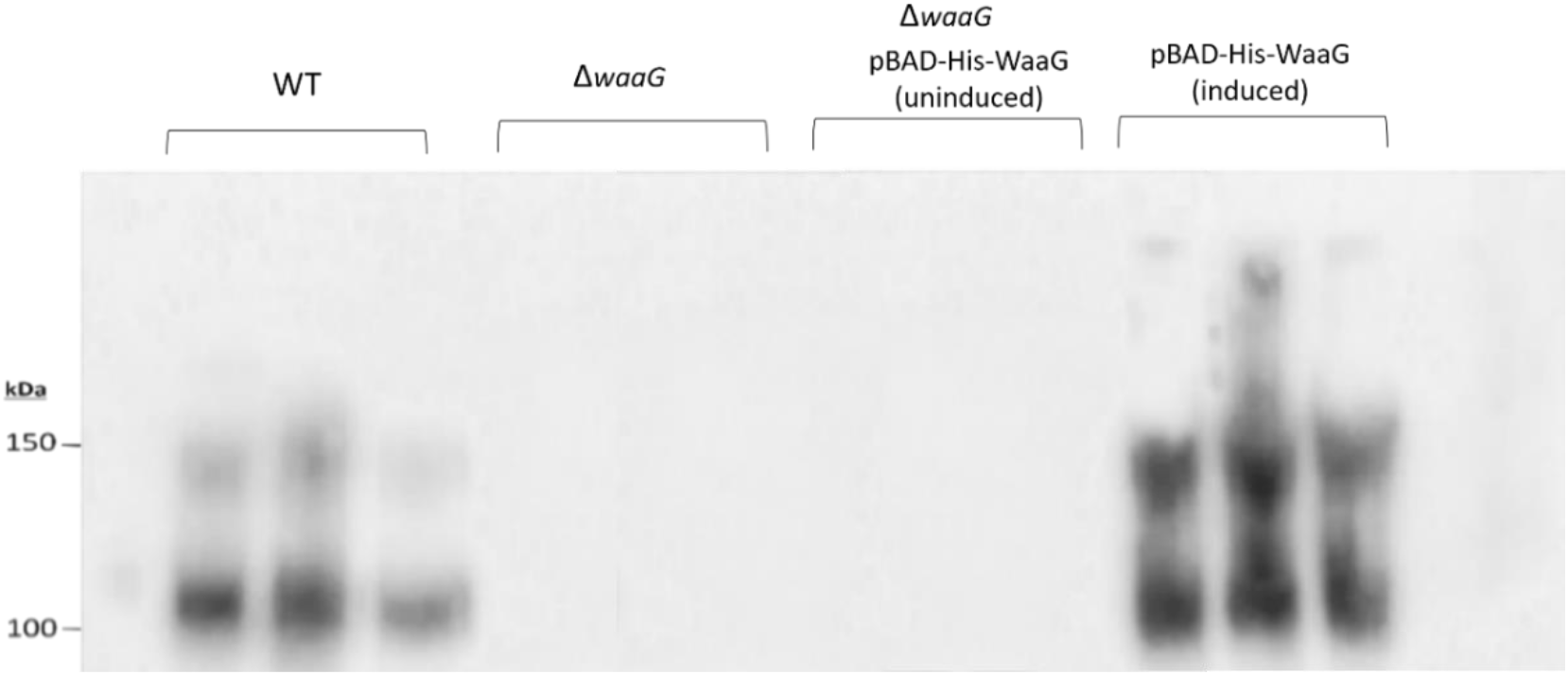
Complementation of Δ*waaG* restores capsule. Western blot analysis of K2 capsule in wild-type, *waaG* mutant and complemented mutant under inducing conditions (0.2% w/v L-arabinose). N=3 biological replicates performed and depicted in the above figure. WT = Wild-type CFT073.

**Figure 7.**
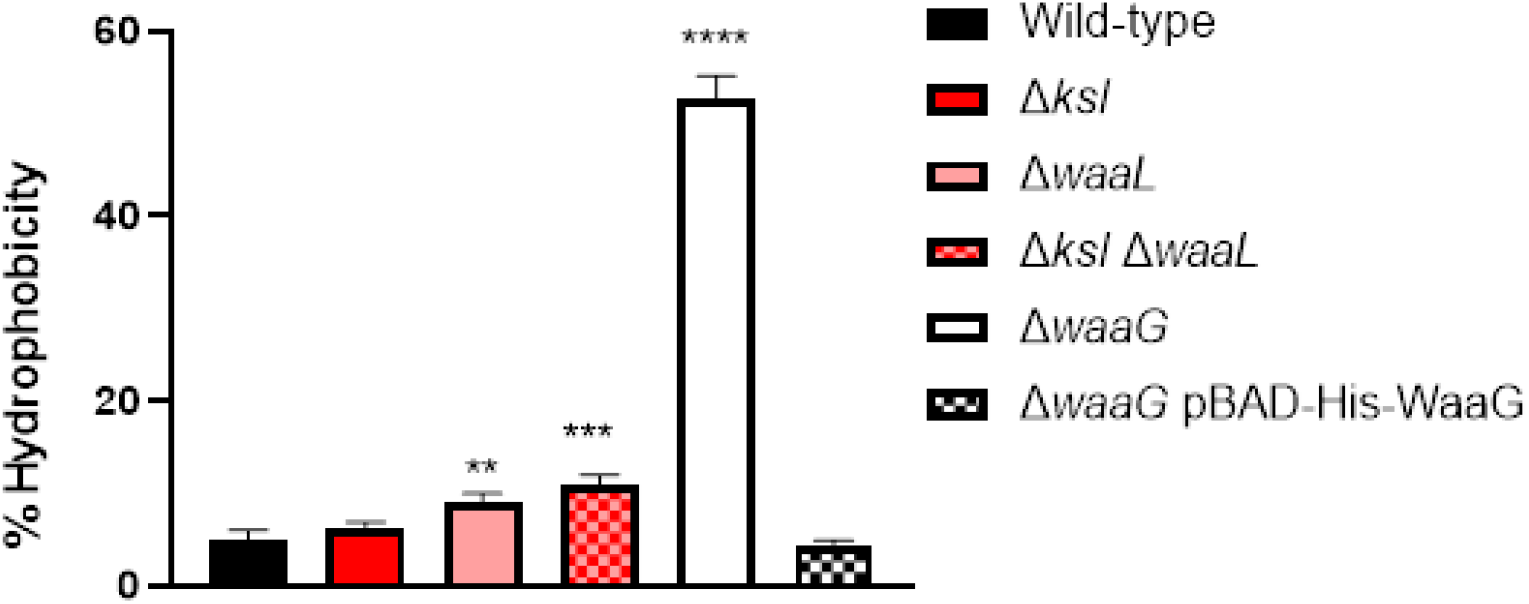
Increased surface hydrophobicity in LPS mutants. Surface hydrophobicity was determined via the n-hexadecane binding assay, as detailed in (37). Percentage hydrophobicity was calculated by dividing the OD_600nm_ of the aqueous phase by the starting OD_600nm_ of 1 (x100), following the incubation of cells with n-hexadecane. N = 3. Statistical significance calculated by One-way ANOVA followed by Dunnett’s multiple comparisons. Wild-type = Wild-type CFT073. ** = P≤0.01, *** = P ≤0.001, **** = P ≤0.0001.

### LPS mutants display changes in surface hydrophobcity

Surface hydrophibicity of the ExPEC derivatives was determined using the n-hexadecane hydrophobicity assay, as detailed in (37). LPS mutants including Δ*waaL*, Δ*waaL* Δ*ksl* and Δ*waaG* displayed significant changes in surface hydrophobicity relative to wild-type CFT073. The outer core mutant deficient in *waaG* displayed the most significant change in surface hydrophobicity, in fact, the strain displayed an approximate 10-fold increase in hydrophobicity (52% cells in aqueous phase) relative to wild-type (5% cells). These results demostrate the significant changes which occur to CFT073 cell surface charge in the absence of LPS, thus affecting capsule retention.

### Outer core mutants possess a punctate K2 expression pattern

Despite the presence of decreased capsule in O6 antigen and outer core mutant derivatives of CFT073, as seen in Figure 2 (Western blot on whole-cell lysates) the mutants do still possess some K2 capsule. In order to discern whether this capsule is intracellular or on the cell surface, immunofluorescence microscopy was performed on exponential and stationary phase CFT073 cultures as depicted in Figure 8. These results showed that at both exponential and stationary phase, *waaG* mutant cells appear more rounded in comparison to the typical rod-shape seen in wild-type cells. Moreover, the *waaG* mutants display a punctate capsule pattern, compared to the homogenous coating of capsule seen on the surface of wild-type cells. Interestingly, despite *waaL* mutants displaying reduced capsule, the immunofluorescence microscopy showed that pattern of capsule association on the cell surface is comparable to that of the wild-type capsule, although with reduced signal. These results indicate changes to the export and localisation of capsule in *waaG* mutants in addition to reduced retention of surface capsule, whilst *waaL* mutants solely display decreased surface capsule without changes to localisation patterns.

**Figure 8.**
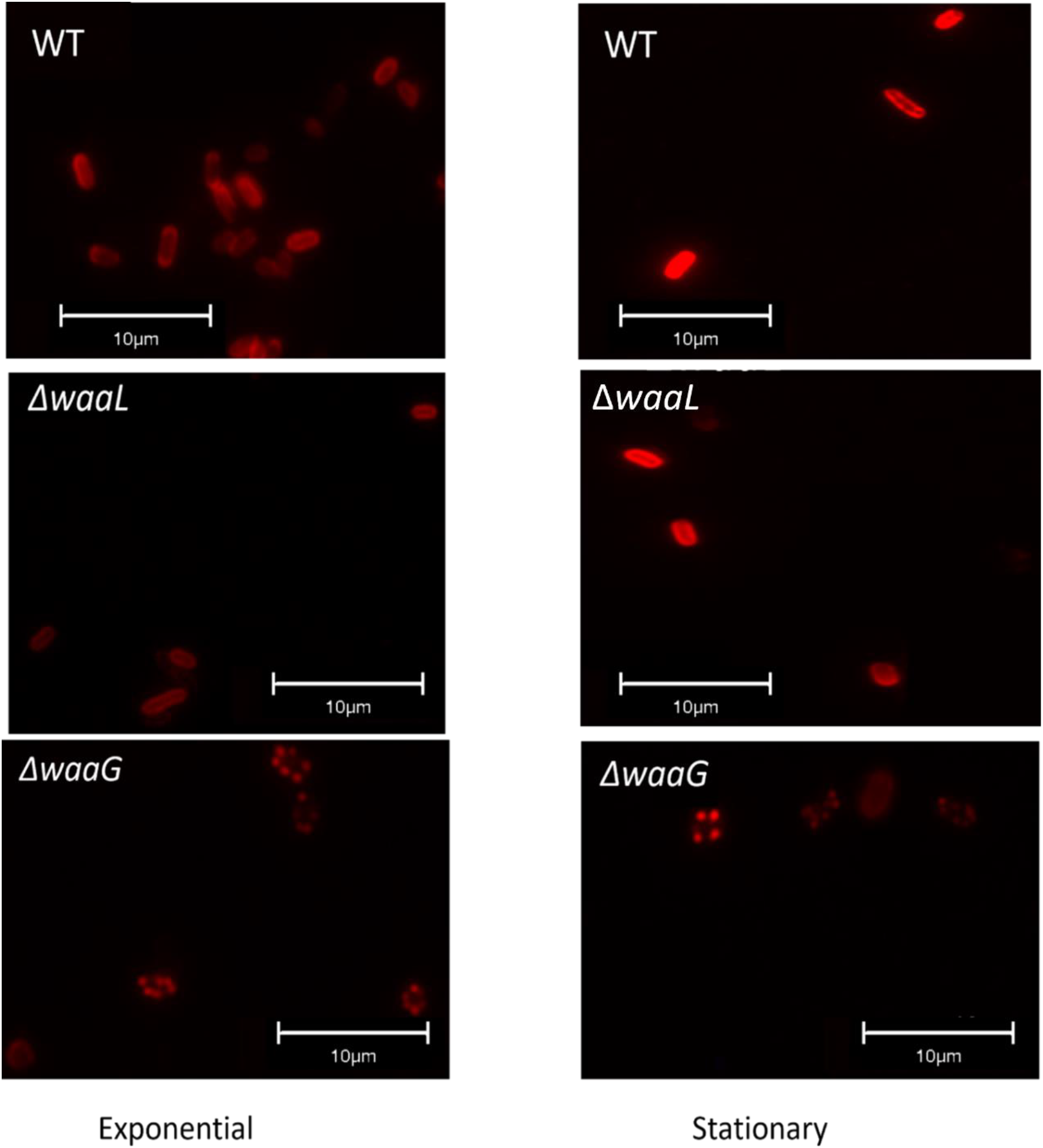
*waaG* mutants display punctate capsule pattern. (A) Exponential and (B) Stationary phase CFT073 cultures were probed with anti-K2 antibody and anti-rabbit conjugated to AlexaFluor594 to detect surface capsule. N = 4 biological replicates. N = 1 shown per growth phase. WT = CFT073 Wild-type.

### Complementation of WaaG restores capsule localisation

Complementation of the *waaG* mutants with pBAD-His-WaaG resulted in restored capsule levels comparable to wild-type as well as full length LPS, as shown in Figure 6, and so it was hypothesised that complemented mutants would also display restored capsule localisation and coverage of the bacterial cell. Immunofluorescence microscopy was carried out on wild-type (not shown) and Δ*waaG* cells complemented with pBAD-His-WaaG under inducing conditions (0.2% L-arabinose - WaaG expression is under regulation of the P*araBAD* promoter), as well as non-inducing conditions as a control. Figure 9A depicts *waaG* mutants complemented with the pBAD-His-WaaG vector under non-inducing conditions (i.e., no L-arabinose added). The cells display the same rounded shape and punctate capsule pattern observed in Figure 8. Interestingly, *waaG* mutants complemented with the pBAD-His-WaaG vector under inducing conditions display wild-type capsule patterns a more defined rod shape, confirming that the absence of outer core and O antigen contribute to the rounding of cells in addition to changes in capsule localisation and surface expression. Additionally, 0.0005% Hoechst 33342 in 80% glycerol was added to microscope slides prior to the addition of the cover slip in order to stain the DNA of bacterial cells to confirm the punctate patterns seen in *waaG* mutants were in-fact cells and not extracellular capsule. Figure 9B depicts that the punctate cells are in fact bacterial cells due to the dual staining of capsule and bacterial DNA.

**Figure 9.**
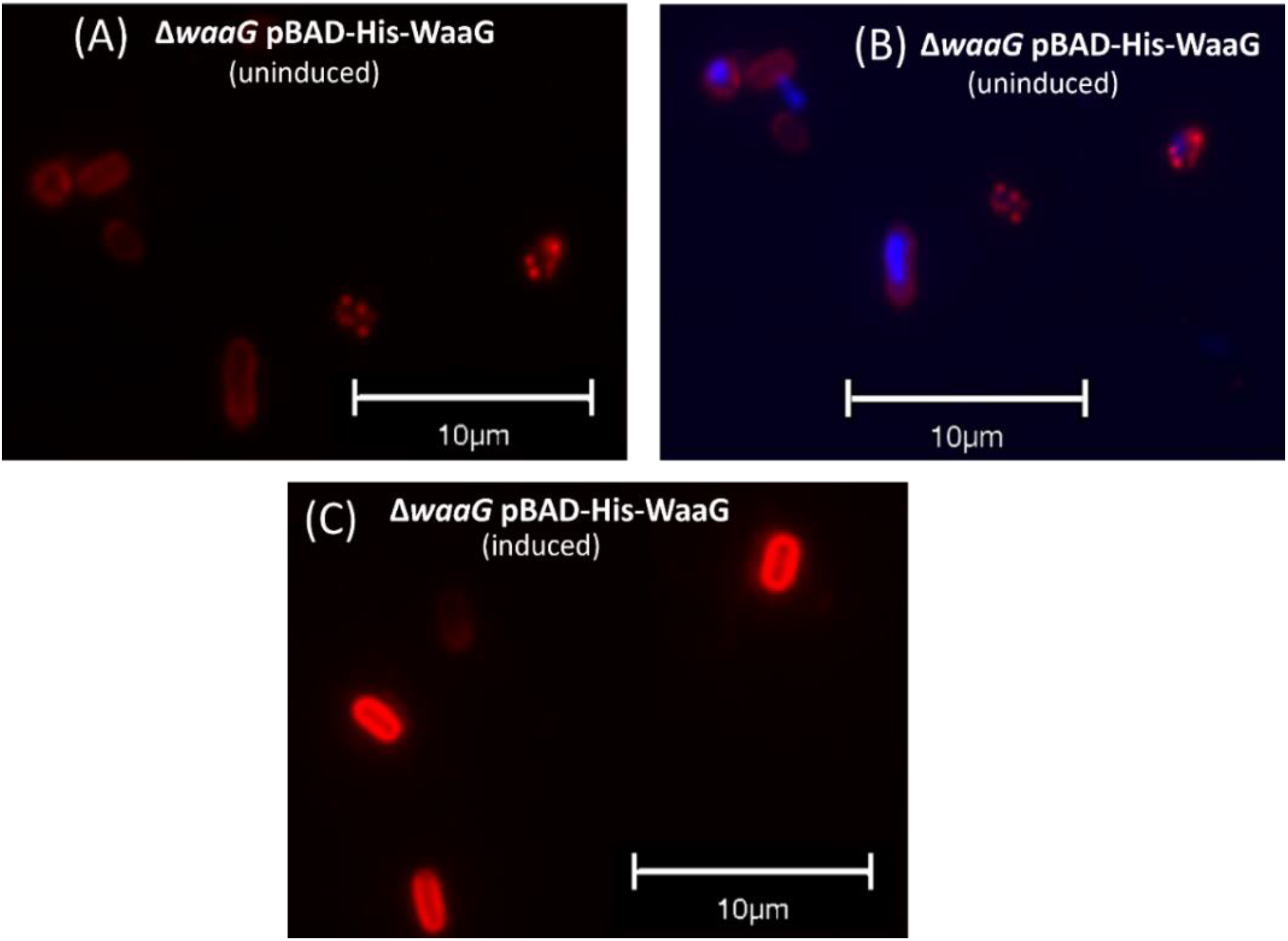
Complemented *waaG* mutants display wild-type capsule localisation and surface expression. Stationary phase cultures of wild-type (not shown) and Δ*waaG* pBAD-His-WaaG grown under non-inducing (A, B) and inducing (C) conditions were probed with anti-K2 antibody and anti-rabbit conjugated to AlexaFluor594 to detect surface capsule. Images in (A) and (C) were taken under a Texas Red filter, whilst (B) depicts an overlay of images taken under Texas Red and DAPI filters. N = 3 biological replicates. N =1 replicate shown. Uninduced = grown in LB. Induced = grown in LB supplemented with 0.2% w/v L-arabinose.

### Contribution of glycome to serum resistance

It has been shown that the K2 capsule and O6 antigen of ExPEC strain CFT073 confer resistance to serum (12, 13, 38). However, a strain CFT073 mutant deficient in both K2 and O6 had not been explored in order to estimate the full contribution of the extracellular glycome to serum resistance. Wild-type was largely resistant to 50% NHS at exponential growth phase (83% survival compared to wild-type cells exposed to HIS). Capsule mutant Δ*ksl* displayed a significant reduction in serum resistance (4% survival) in comparison to wild-type (*p* <0.0001) during exponential growth. Similarly, O antigen deficient Δ*waaL* in exponential phase displayed significantly reduced survival in 50% NHS (0.01% survival) in comparison to the wild-type (*p* <0.0001). Strikingly, the double mutant Δ*ksl* Δ*waaL* was almost entirely sensitive to 50% NHS during exponential growth phase, with only 0.000038 % of cells surviving. This reduction in survival was statistically significant when compared with the percentage survival of wild-type CFT073 (*p* < 0.0001). Only 0.009% of exponential *waaG* survived when exposed to 50% NHS, a significant reduction from wild-type survival (*p* <0.0001). Furthermore, the survival of stationary phase CFT073 strains after exposure to 50% NHS for 90 minutes was examined. Stationary phase wild-type cultures displayed resistance to 50% NHS with 90% of cells surviving. The slight reduction in survival of exponential (83%) in comparison to stationary phase (90%) wild-type CFT073 was not significant (*p* = 0.4). Stationary phase Δ*ksl* and Δ*waaL* mutants were significantly more sensitive to serum than wild-type, with 4% and 0.2% survival, respectively (*p* < 0.0001 for both mutants relative to wild-type). Moreover, the double mutant, Δ*ksl* Δ*waaL*, was again, exquisitely sensitive to 50% NHS during stationary phase (0.0005% survival, *p* < 0.0001, relative to wild-type). 0.1% of *waaG* mutants survived in 50% NHS at stationary phase growth, which was a significant reduction compared to wild-type (*p* <0.0001). Interestingly, there was a significant increase in the survival of *waaL* mutants during stationary phase compared with exponential growth, *p* = 0.0346. Overall, these data indicate that LPS and capsule are the major determinants of serum resistance, as a double mutant deficient in O antigen and capsule synthesis is almost entirely sensitive to serum, with no significant difference in survival during exponential and stationary phase serum resistance between this mutant and a K-12 serum sensitive control, DH5α (data not shown as DH5α cultures display 0% survival after exposure to 50% NHS for 90 minutes).

### WaaG inhibition alters LPS electrophoretic mobility and reduces surface K2 capsule

Published works by Muheim *et al* identified a WaaG inhibitor with *in vitro* activity towards purified LPS, 4-(2-amino-1,3-thiazol-4-yl) (known as L1) with IC_50_ = 1.0 mM (39). Due to the role for WaaG in capsule retention identified in this study (by virtue of its contribution to LPS composition and surface charge), it was hypothesised that L1 could be used therapeutically to reduce surface association of K2 capsule with the cell and increase the sensitivity of CFT073 to antibiotics. Moreover, the use of L1 or other LPS inhibitors to reduce CFT073 virulence and antibiotic susceptibility is of additional benefit as the compound is non-toxic to bacteria, meaning L1 or a derivative could be used therapeutically and without damage to the microbiome. Concentrations of L1 at 1.25 mM and 3.125 mM were explored for the inhibition of WaaG in live bacterial cultures of CFT073. L1 was solubilised in DMSO and so cultures with added DMSO, along with the *waaG* mutant were used as a control. As seen in Figure 10, the addition of L1 to cultures results in a concentration-dependent reduction in capsule (A), in addition to changes in LPS quantity and electrophoretic mobility (B). Taken together, the above results show that L1 can reduce capsule levels through LPS inhibition, and also displays its inhibitory activity on live bacterial cells and not just purified LPS, as had been explored in (39).

**Figure 10.**
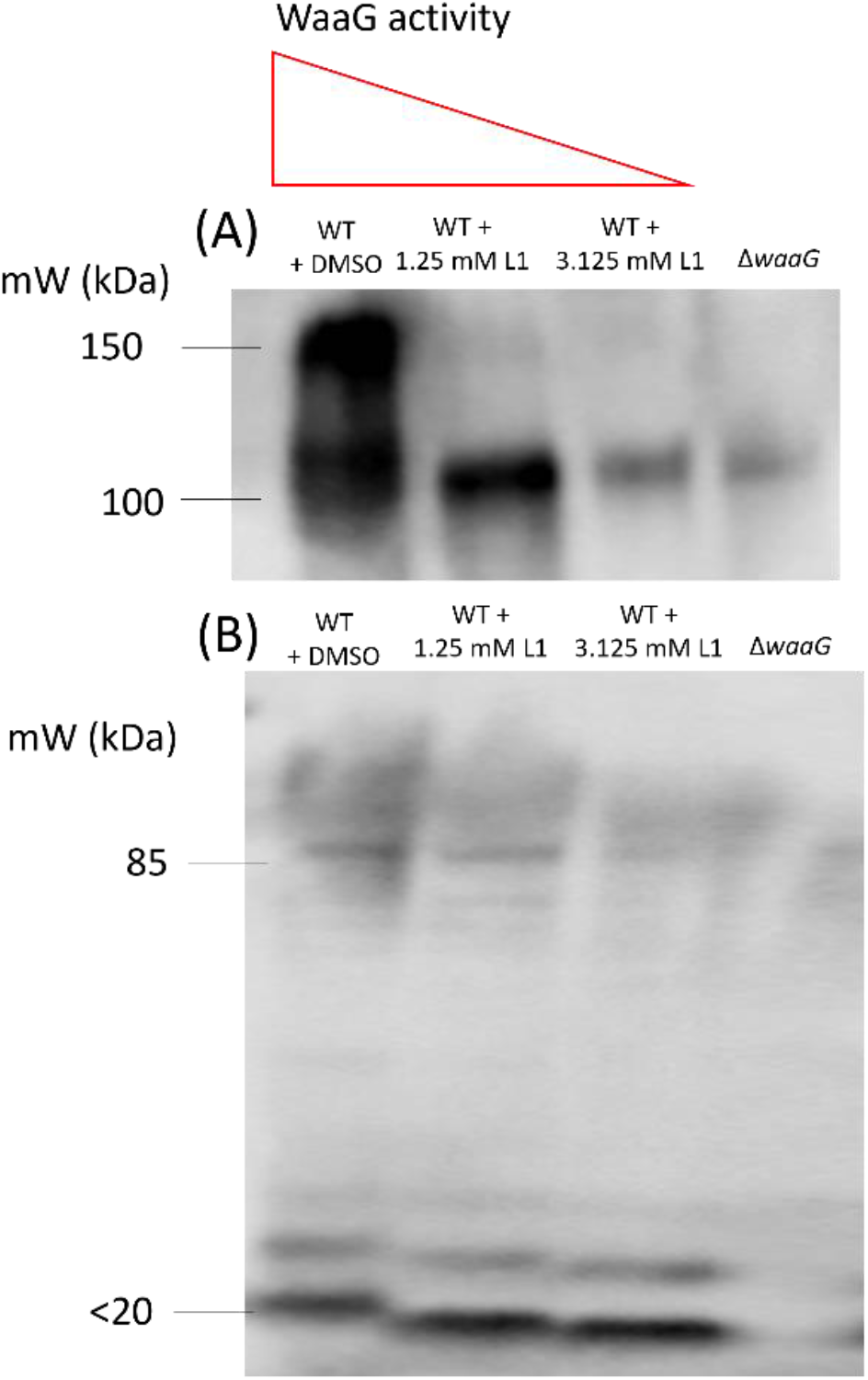
WaaG inhibition reduces K2 capsule. Whole cell lysates of the WaaG deficient mutant Δ*waaG* and Wild-type CFT073 supplemented with DMSO or L1 at concentrations of 1.25 or 3.125 mM were separated by SDS-PAGE and probed with (A) anti-K2 and (B) anti-O6 antibodies for Western blot analysis of LPS and capsule. N=3 conducted, N=1 shown.

### LPS/capsule synergy can be exploited to reduce serum resistance

The reduced K2 capsule levels in cultures exposed to the WaaG inhibitor L1 were quantified by (A) densitometry performed on Western blots and (B) ELISA and compared to the wild-type (+DMSO) and the *waaG* mutant. Figure 11A depicts densitometric quantification of K2 capsule in N = 4 Western blots, demonstrating that wild-type + L1 at 1.25 mM and 3.125 mM display significant reductions to capsule levels, compared to wild-type + DMSO (*p* < 0.0001 for both). Interestingly, the addition of 3.125 mM L1 reduced capsule to just 17% of wild-type levels, which was comparable to the amount of K2 displayed by the *waaG* mutant (*p* > 0.9999). ELISA quantification of capsule (Figure 11B) mirrored those seen in the densitometric analysis, where the addition of L1 reduced K2 levels when compared to wild-type + DMSO in a concentration-dependent manner (1.25 mM L1; *p* = 0.07, 3.125 mM L1; *p* = 0.0009). Finally, it was determined whether the changes to the glycome mediated by L1 would alter the resistance of CFT073 to 50% human serum (NHS). As shown in Figure 11C, the addition of L1 resulted in significant reductions to CFT073 serum survival in a concentration-dependent manner (*p* ≤ 0.00001 for wild-type + L1 at 1.25 and 3.125 mM, as well was the *waaG* mutant) when compared to the wild-type + DMSO.

**Figure 11.**
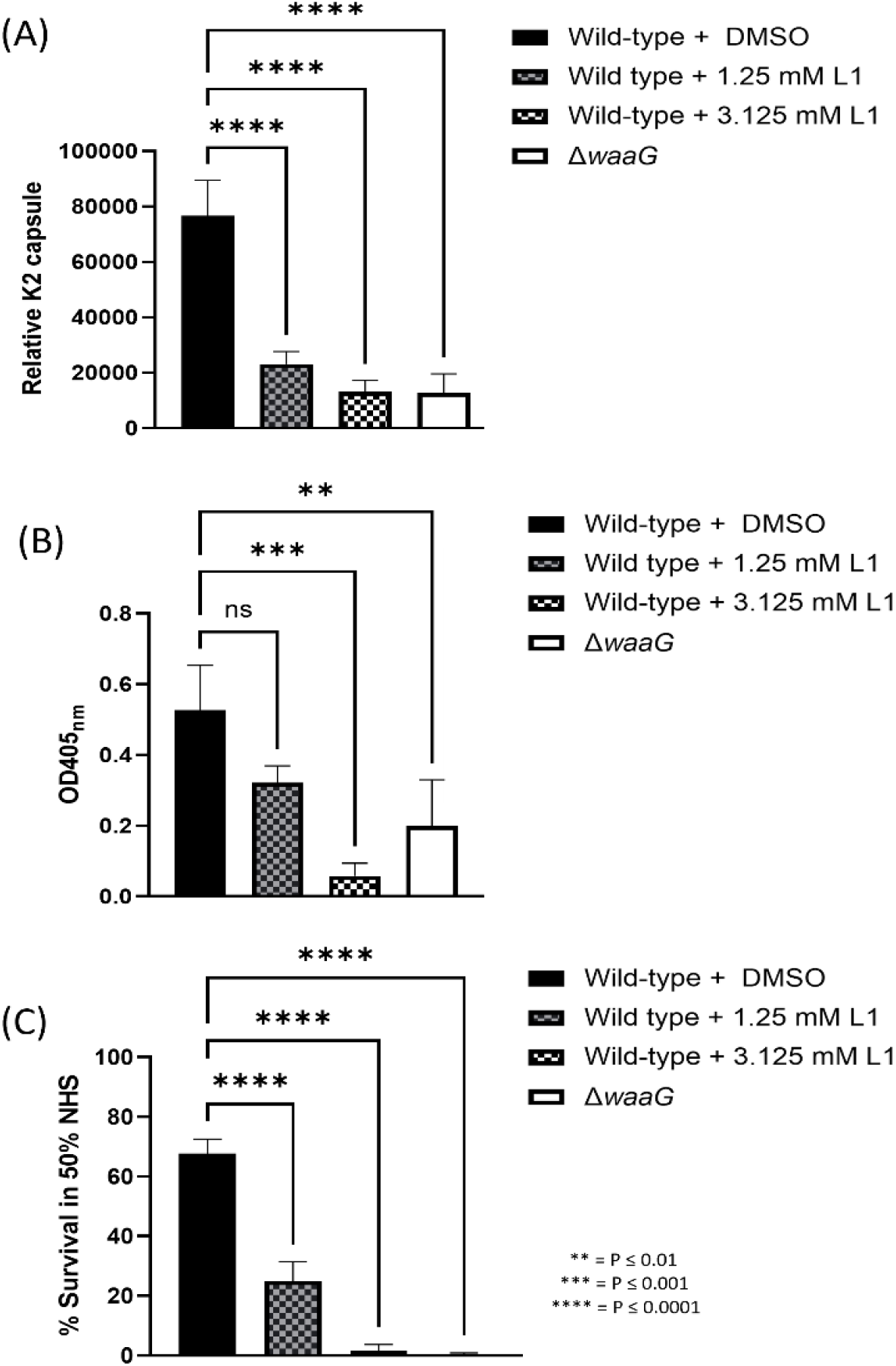
Implications of WaaG inhibition on capsule levels and serum resistance. (A) Densitometry performed on anti-K2 Western blots, N = 3. (B) ELISA examining K2 levels conducted on Wild-type CFT073 +/−the addition of L1, in addition to the *waaG* mutant, N=3. (C) Serum killing assay, % survival calculated as a fraction of CFU/mL at T90 in NHS over CFU/mL at T90 in HIS (x100). N = 4. Statistical significance calculated by One-way ANOVA and Dunnett’s multiple comparisons.

### WaaG inhibition increases sensitivity of CFT073 to rifampicin

Lastly, the impact of L1-dependent WaaG inhibition on antibiotic susceptibility was explored. It is known that mutations to genes implicated in LPS core biosynthesis confer significant changes to cell permeability, in addition to displaying increased susceptibility to hydrophobic compounds and antibiotics (40). Cultures grown with or without the addition of L1 (in addition to the *waaG* mutant) were exposed to varying concentrations of rifampicin, as shown in Table 2. The addition of 1.25 mM and 3.125 mM of L1 significantly reduced the minimum inhibitory concentration (MIC) of CFT073 to rifampicin, compared to the wild-type + DMSO. As expected, the *waaG* mutant also displayed a significant increase in sensitivity to rifampicin (*p* ≤ 0.00001).

**Table 1.** Extracellular glycome mutants are sensitive to 50% NHS. Percentage survival was calculated as a fraction of CFU/mL at T90 in NHS over CFU/mL at T90 in HIS (x100). N = 4 for both assays. WT = Wild-type CFT073. Statistical significance calculated by Two-way ANOVA and Tukey’s multiple comparisons.

| Strain | Exponential | Stationary |
| --- | --- | --- |
| Wild-type | 85 | 90 |
| $\Delta ksl$ | 4 * | 4 * |
| $\Delta waaL$ | 0.008 ** | 0.19 ** |
| $\Delta ksl \Delta waaL$ | 0.00004 ** | 0.0005 ** |
| $\Delta waaG$ | 0.0006% ** | 0.1125% ** |

**Table 2.** WaaG inhibitor L1 reduces CFT073 sensitivity to rifampicin. N = 3. Statistical significance calculated by One-way ANOVA and Dunnett’s multiple comparisons.

| Strain | Average MIC (N=3) | Significance (relative to wild-type) |
| --- | --- | --- |
| Wild-type + DMSO | 12.5 µg/ mL |  |
| Wild-type + 1.25 mM L1 | 10 µg/ mL | ** |
| Wild-type + 3.125 mM L1 | 2 µg/ mL | **** |
| $\Delta waaG$ | 2 µg/ mL | **** |

## Discussion

### Visualisation of CFT073 glycome

To survive in serum, bacteria must evade the host immune system including the components of the complement system. Resistance to killing by serum is achieved (in part) by the extracellular glycome of polysaccharides such as LPS, colanic acid and capsule, which act as steric barriers to the complement system, the preliminary obstacle faced by ExPEC upon entry into the bloodstream (12, 13). To further characterise the role of the extracellular glycome in CFT073 serum resistance, a panel of mutants deficient in extracellular polysaccharides O6 antigen, capsule and the outer core oligosaccharide were made. The extracellular glycome profiles of CFT073 wild-type and mutant derivatives were visualised and analysed using techniques such as SDS-PAGE, Western blot, and ELISA. As depicted in Figure 1D, LPS staining of whole cell lysates separated by SDS-PAGE allows for the visualisation of the extracellular glycome components O6 antigen and K2 capsule simultaneously, which marks the first co-visualisation of these serotype-specific polysaccharides on an SDS-PAGE gel. Moreover, the stained SDS-PAGE gels depicted the larger ‘molecular weight’ of K2 in comparison to O6 antigen, with approximately N=40 units to N=25 for the O6 antigen. The results of the SDS-PAGE gels were reproducible, including when conducted at different growth stages, indicating the length and size of these polysaccharides are not phase variable. Western blot analyses of the K2 capsule (such as that show in Figure 2B) depicted 2 bands which resolve at approximately 100 and 150 kDa, an observation previously made in (32), which was also reproducible regardless of cell concentration or growth phase.

### Role of extracellular glycome in serum resistance

The relative survival in 50% NHS of CFT073 and glycome mutants at exponential and stationary phase was examined in this study. During exponential and stationary phase growth, both CFT073 Δ*ksl* and CFT073 Δ*waaL* displayed reduced survival rates in serum in comparison to wild-type CFT073. Interestingly, the *waaL* and *waaG* mutants were more sensitive to serum than the *ksl* mutant at both growth phases, although this is likely due to the fact that CFT073 LPS mutants also display significant reductions in surface K2 capsule. Interestingly, the double O antigen/capsule mutant, Δ*ksl* Δ*waaL* displayed the highest drop in survival relative to wild-type, with the CFU/mL counts of 2/4 exponential biological replicates being reduced to 0, indicating complete killing of the strain by complement. These data reinforce that the O antigen and capsule are the two main determinants of serum resistance in ExPEC strain CFT073, and that exploiting their synergy is a potential therapeutic strategy.

### Synergy between the R1 core, O6 antigen and the K2 capsule

The anti-K2 Western blots conducted on whole cell lysates depicted an apparent reduction in K2 capsule in *waaL* and *waaG* mutants, compared to wild-type CFT073. This reduction was quantified through densitometry and ELISA (Figure 3A and B). Interestingly, CFT073 *waaL* mutants displayed on average a 50% reduction to capsule levels, whilst *waaG* mutants displayed only 10-12% of wild-type capsule levels, as determined by densitometry. These results imply that both O antigen and the LPS core are interacting with K2 to contribute to its retention at the cell surface. Mutants deficient in *rfaH* and *wzy* also displayed reductions to K2 capsule, further reinforcing that the interactions between LPS and capsule retain K2 at the surface and the genes are not involved in cross-regulation or a feedback loop (data not shown).

The reliance of capsule on LPS core moieties, has previously been observed in *E. coli* but only in a K1:O18 background in strain RS218 (33). Thus, the findings of this chapter confirm that the interactions between LPS and capsule are not just K1 and K5 specific, but also occur in K2 *E. coli* and likely other K types. Similarly, in *Klebsiella pneumoniae*, an intact LPS core molecule has been shown to contribute to capsule retention for several different O and K serotypes in strains 52145 (O1:K2), DL1 (O1:K1) and C3 (O8:K66) (35). However, in both of the aforementioned studies (*E. coli* K1 and *Klebsiella*), *waaL* mutants did not display reduced capsule relative to wild-type *E. coli* and *Klebsiella pneumoniae*, implicating solely the charge of the core molecule in capsule retention for these prototypic strains (33, 35). Perhaps the interaction of O6 with K2 in CFT073 is strain-specific or serotype-specific, whilst the core-capsule interaction is more conserved across different strains and serotypes. Supporting this hypothesis is a study conducted on a different *Klebsiella pneumoniae* strain to those studied in (35), in which *Klebsiella pneumoniae* B5055 (O1:K2) *waaL* mutants displayed a 50% reduction to surface capsule levels (34). As different core types and capsule types will be composed of different sugar structures with different modifications (such as phosphorylation) and subsequently different surface charge, it is not unexpected that whilst the synergy between LPS and capsule is conserved, the specific interactions involved in the surface retention (e.g., in CFT073 K2/ core as well as K2/O6 interact vs in RS218 solely K1/core) differ from strain to strain depending on the serotype.

The R1 core molecule (shared between CFT073 and RS218 – as well as most other ExPEC isolates) is highly phosphorylated, giving rise to the net-negative charge of the core (16). Due to this negative charge, the association of divalent cations such as Mg^2+^ and Ca^2+^ cross-link LPS, provide membrane stability, as well as serve as a Ca^2+^ reservoir for RTX family toxins like HlyA (41). RS218 (K1:O18) mutants deficient in the enzymes catalysing the phosphorylation of the R1 core (such as *waaY -* unphosphorylated HepII) display no structural changes to LPS other than changes to its net charge due to the lack of phosphate groups, yet display only 25% surface capsule compared to wild-type RS218 (33). These results validate the hypothesis that the interactions between LPS and capsule are ionic, physiochemical interactions which result in capsule retention. Additionally, this study has shown that LPS mutant derivatives, particularly *waaG* mutants, display significant changes to surface hydrophobicity which further implicates the importance of electrostatic surface interactions in capsule retention.

In this study, despite the clear reduction in K2 seen in CFT073 *waaL* and *waaG* mutants upon densitometric analysis of Western blots conducted on whole cell lysates (Figure 3A), ELISA data demonstrated a less profound change to capsule levels at both exponential and stationary phase (Figure 3B). Based on the findings from the *E. coli* K1 paper (33), and the findings depicted in Figure 5 which demonstrated that the changes to capsule in LPS mutants were post-transcriptional, it was hypothesised that the capsule not retained on the cell surface was present in the supernatant, thus resulting in higher signals for K2 capsule in the ELISA. The Western blot analyses were conducted on supernatant-free whole cell lysates, thus serving as a more realistic depiction of surface capsule compared to ELISA.

Western blot and ELISA on CFT073 culture supernatants confirmed that (a) under normal conditions there is some K2 capsule secreted into the supernatant by wild-type CFT073 and that (b) *waaL* and *waaG* mutants display significantly more K2 capsule in the supernatant compared to wild-type due to the inability of these mutants to retain capsule by virtue of ionic interactions with the outer core and O antigen (Figure 4A and B). The presence of K2 capsule in the supernatant of wild-type CFT073 cultures can be explained by the various potential roles for free K2 capsule. The CFT073 K2 capsule has been shown to hinder biofilm formation of pathogenic competitor species such as *Pseudomonas aeruginosa, Klebsiella pneumoniae, Staphylococcus aureus, Staphylococcus epidermidis* and *Enterococcus faecalis* (25). These findings provide an explanation for the presence of K2 capsule in wild-type CFT073 supernatant, with changes to LPS giving rise to an increased loss of capsule.

### Localisation of K2 capsule on the cell surface

Following the observation that *waaL* and *waaG* mutants fail to retain K2 capsule at the cell surface, resulting in increased K2 in the supernatant, surface K2 was visualised by immunofluorescence microscopy, as shown in Figure 8. Interestingly, the K2 pattern on wild-type cells at exponential and stationary phase appeared, for some cells, to show more K2 capsule at the cell poles, in comparison to the equator. Other cells showed a more even distribution of K2 capsule. For all wild-type cells, however, surface capsule covered the entire cell. The observation that the cell poles possess more K2 is not unique to CFT073 and the K2 capsule, a study conducted on ExPEC strain EV136 (K1 capsule) showed that capsule export occurs evenly and randomly across the entire surface of the cell, however, the K1 located at the poles was thicker than the equatorial K1 capsule (42). Additionally, the study found that the EV136 K1 capsule displays two qualitatively different domains of bacterial capsule height/length at the poles and at the equator, with the poles possessing longer capsule and the equators possessing shorter polysaccharide chains (42). Similarly, purified K5 capsule possesses two components, denoted ‘high’ and ‘low’ molecular weight components (43). These results provide reason for the difference in signal at the poles in some of the wild-type immunofluorescence microscopy images shown in Figure 8, yet also provide an explanation for the two bands at approximately 100 kDa and 150 kDa visualised upon Western blot analysis of the capsule (such as in Figure 2B).

### Exploitation of LPS/capsule synergy has potential therapeutic benefit

Muheim *et al* showed that an aminothiazole molecule 4-(2-amino-1,3-thiazol-4-yl)phenol inhibits WaaG glycosyltransferase activity *in vitro*, preventing the outer core assembly and thus, O antigen attachment of purified LPS. This study showed that the inhibitor was effective in preventing some of the WaaG activity in CFT073, thereby affecting LPS quantity, integrity and subsequently K2 capsule association. Additionally, L1 treatment of CFT073 resulted in increased sensitivity to a hydrophobic antibiotic, rifampicin, as well as significantly reduced resistance to 50% human serum. Thus, this compound or a derivative with higher affinity for *E. coli* WaaG could be considered as an alternative treatment to antibiotics as inhibition of WaaG, as we have shown, will reduce virulence significantly through preventing O antigen and K2 capsule association with the cell (33). Additionally, the compound (or a similar one) could be co-administered with antibiotics, as truncated LPS will result in increased sensitivity to antibiotics due to altered cell permeability and changes to surface charge (40). Moreover, alternative therapies such as phage therapy could be considered, as many phages target the extracellular glycome, and several K/O antigens are correlated specifically with ExPEC infections (44, 45). For example, O antigen depolymerases could be isolated and cloned from phage for therapeutic use, which would also result in reduced capsule and overall virulence (46). Targeting the glycome through phage therapy or WaaG inhibitors will do little to affect the microbiome and so again, are two promising avenues of research.

## Conclusions

The synergy between LPS and capsule observed in *E. coli* K1, K5 and several serotypes of *Klebsiella pneumoniae* also exists in strain CFT073 which possesses the R1 core, O6 antigen and K2 capsule. Due to the existence of the interactions between LPS and capsule across other species and strains, the synergy may be conserved across other members of the *Enterobacteriales* and requires exploration. This synergy can be therapeutically targeted in order to sensitise bacteria to antibiotics and reduce overall virulence. The CFT073-specific interactions between components of the extracellular glycome is graphically summarised in Figure 12.

**Figure 12.**
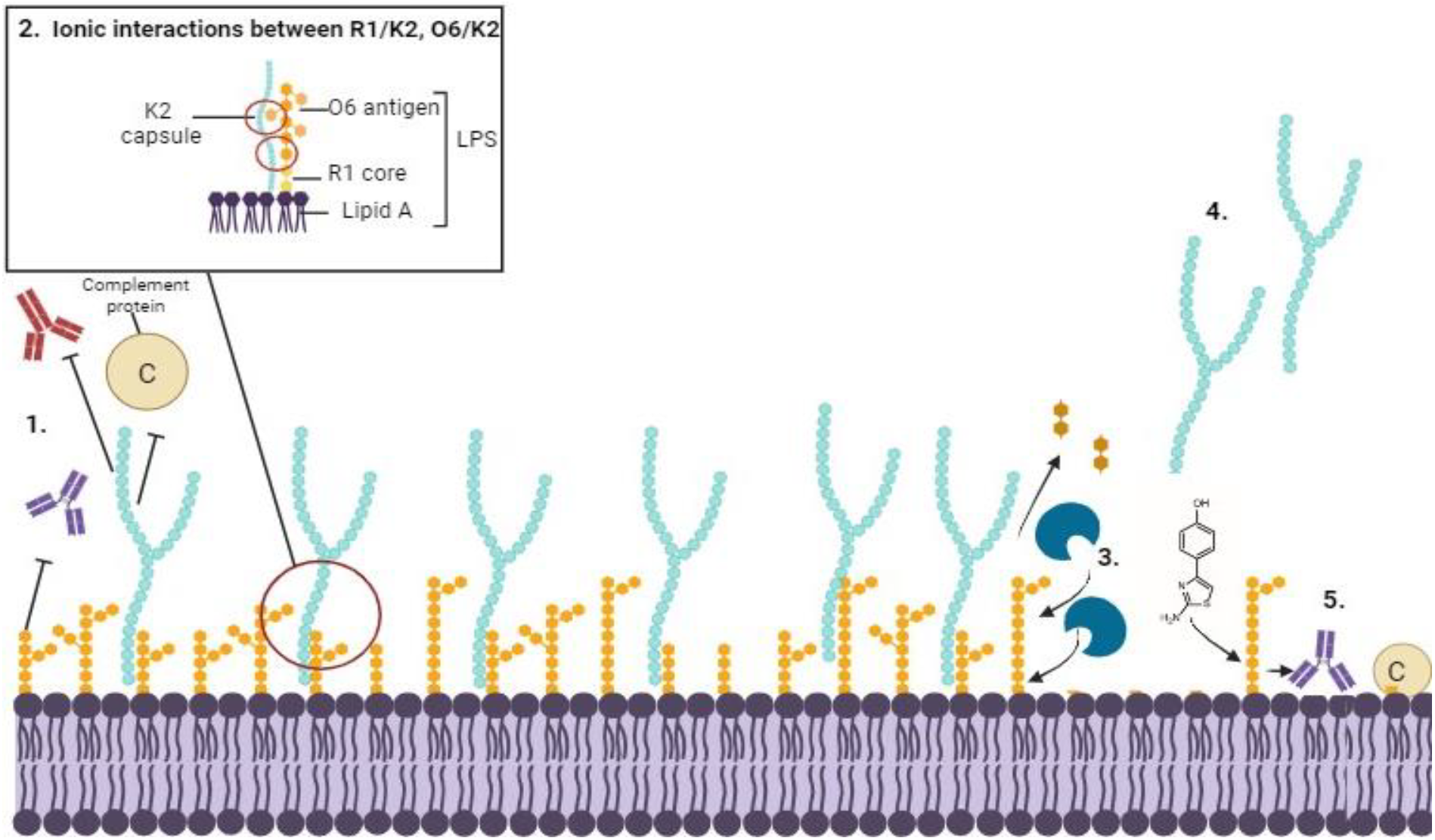
Summary of interactions between LPS and K2 and implications. Graphical depiction of the CFT073 outer leaflet, including extracellular polysaccharides LPS and K2. (1) Under normal conditions, LPS O antigen and the K capsule provide steric barriers to antibodies and the complement system, enabling disseminated infection and sepsis. (2) The K2 capsule is retained at the cell surface through interactions with the O6 antigen and the R1 outer core, as well as Braun’s Lpp (not shown). (3) LPS-capsule synergy can be exploited through treatment with LPS depolymerases (blue) or WaaG inhibitors such as L1. (4) Depolymerisation of LPS or WaaG inhibition result in increased cell permeability and reduced capsule association, allowing (5) access to the bacterial cell surface for antibody binding and complement protein deposition, in addition to increased antibiotic sensitivity.

## Author statements

## 1.6 Author contributions

N.McG and S.S conceived of and designed the experiments. N.McG performed all experiments bar those depicted in Figure 11, which were performed by D.R. N.McG and S.S analysed all of the data.

D.R analysed the data pertaining to experiments depicted in Figure 11. N.McG wrote the article.

## 1.9 Ethical approval

*N/A*

## 1.10 Consent for publication

N/A

## 1.11 Acknowledgements

Plasmid pBAD-His-WaaG was kindly supplied by Daniel Daley. N McG and SS acknowledge DR for his assistance with the WaaG inhibitor experiments.

## References

1. Vihta KD, Stoesser N, Llewelyn MJ, Quan TP, Davies T, Fawcett NJ, et al. Trends over time in Escherichia coli bloodstream infections, urinary tract infections, and antibiotic susceptibilities in Oxfordshire, UK, 1998-2016: a study of electronic health records. Lancet Infect Dis. 2018;18(10):1138–49.

2. Foxman B. The epidemiology of urinary tract infection. Nat Rev Urol. 2010;7(12):653–60.

3. Ciesielczuk H, Jenkins C, Chattaway M, Doumith M, Hope R, Woodford N, et al. Trends in ExPEC serogroups in the UK and their significance. Eur J Clin Microbiol Infect Dis. 2016;35(10):1661–6.

4. Phan MD, Peters KM, Sarkar S, Lukowski SW, Allsopp LP, Gomes Moriel D, et al. The serum resistome of a globally disseminated multidrug resistant uropathogenic Escherichia coli clone. PLoS Genet. 2013;9(10):e1003834.

5. Hertz FB, Nielsen JB, Schønning K, Littauer P, Knudsen JD, Løbner-Olesen A, et al. “Population structure of drug-susceptible,-resistant and ESBL-producing Escherichia coli from community-acquired urinary tract”. BMC Microbiol. 2016;16:63.

6. Miajlovic H, Aogáin MM, Collins CJ, Rogers TR, Smith SG. Characterization of Escherichia coli bloodstream isolates associated with mortality. J Med Microbiol. 2016;65(1):71–9.

7. Li D, Elankumaran P, Kudinha T, Kidsley AK, Trott DJ, Jarocki VM, et al. Dominance of. Microb Genom. 2023;9(7).

8. Dale AP, Woodford N. Extra-intestinal pathogenic Escherichia coli (ExPEC): Disease, carriage and clones. J Infect. 2015;71(6):615–26.

9. Miajlovic H, Smith SG. Bacterial self-defence: how Escherichia coli evades serum killing. FEMS Microbiol Lett. 2014;354(1):1–9.

10. Fratamico PM, DebRoy C, Liu Y, Needleman DS, Baranzoni GM, Feng P. Advances in Molecular Serotyping and Subtyping of Escherichia coli. Front Microbiol. 2016;7:644.

11. Miajlovic H, Cooke NM, Moran GP, Rogers TR, Smith SG. Response of extraintestinal pathogenic Escherichia coli to human serum reveals a protective role for Rcs-regulated exopolysaccharide colanic acid. Infect Immun. 2014;82(1):298–305.

12. Sarkar S, Ulett GC, Totsika M, Phan MD, Schembri MA. Role of capsule and O antigen in the virulence of uropathogenic Escherichia coli. PLoS One. 2014;9(4):e94786.

13. Buckles EL, Wang X, Lane MC, Lockatell CV, Johnson DE, Rasko DA, et al. Role of the K2 capsule in Escherichia coli urinary tract infection and serum resistance. J Infect Dis. 2009;199(11):1689–97.

14. Kalynych S, Morona R, Cygler M. Progress in understanding the assembly process of bacterial O-antigen. FEMS Microbiol Rev. 2014;38(5):1048–65.

15. McCabe WR, Kaijser B, Olling S, Uwaydah M, Hanson LA. Escherichia coli in bacteremia: K and O antigens and serum sensitivity of strains from adults and neonates. J Infect Dis. 1978;138(1):33–41.

16. Amor K, Heinrichs DE, Frirdich E, Ziebell K, Johnson RP, Whitfield C. Distribution of core oligosaccharide types in lipopolysaccharides from Escherichia coli. Infect Immun. 2000;68(3):1116–24.

17. Dissanayake DR, Wijewardana TG, Gunawardena GA, Poxton IR. Distribution of lipopolysaccharide core types among avian pathogenic Escherichia coli in relation to the major phylogenetic groups. Vet Microbiol. 2008;132(3-4):355–63.

18. Leclercq SO, Branger M, Smith DGE, Germon P. Lipopolysaccharide core type diversity in the. Microb Genom. 2021;7(9).

19. Chakraborty A, Saralaya V, Adhikari P, Shenoy S, Baliga S, Hegde A. Characterization of Escherichia coli Phylogenetic Groups Associated with Extraintestinal Infections in South Indian Population. Ann Med Health Sci Res. 2015;5(4):241–6.

20. Kim KS, Itabashi H, Gemski P, Sadoff J, Warren RL, Cross AS. The K1 capsule is the critical determinant in the development of Escherichia coli meningitis in the rat. J Clin Invest. 1992;90(3):897–905.

21. van Alphen L, Arends A, Hopman C. Cross-reactions between Neisseria meningitidis group H and Escherichia coli K2 and K62 polysaccharides. J Clin Microbiol. 1988;26(2):394–6.

22. Hirschel B, Boulnois G, Timmis K. [Neisseria meningitidis B and E. coli K1: different genes causing the production of an identical capsule. Preliminary communication]. Schweiz Med Wochenschr. 1983;113(49):1857–8.

23. Whitfield C. Biosynthesis and assembly of capsular polysaccharides in Escherichia coli. Annu Rev Biochem. 2006;75:39–68.

24. Whitfield C. Structure and Assembly of Escherichia coli Capsules. EcoSal Plus. 2009;3(2).

25. Valle J, Da Re S, Henry N, Fontaine T, Balestrino D, Latour-Lambert P, et al. Broad-spectrum biofilm inhibition by a secreted bacterial polysaccharide. Proc Natl Acad Sci U S A. 2006;103(33):12558–63.

26. Goller CC, Seed PC. Revisiting the Escherichia coli polysaccharide capsule as a virulence factor during urinary tract infection: contribution to intracellular biofilm development. Virulence. 2010;1(4):333–7.

27. Lloyd AL, Henderson TA, Vigil PD, Mobley HL. Genomic islands of uropathogenic Escherichia coli contribute to virulence. J Bacteriol. 2009;191(11):3469–81.

28. Devine DA, Roberts AP. K1, K5 and O antigens of Escherichia coli in relation to serum killing via the classical and alternative complement pathways. J Med Microbiol. 1994;41(2):139–44.

29. Suerbaum S, Friedrich S, Leying H, Opferkuch W. Expression of capsular polysaccharide determines serum resistance in Escherichia coli K92. Zentralbl Bakteriol. 1994;281(2):146–57.

30. Rigg GP, Barrett B, Roberts IS. The localization of KpsC, S and T, and KfiA, C and D proteins involved in the biosynthesis of the Escherichia coli K5 capsular polysaccharide: evidence for a membrane-bound complex. Microbiology (Reading). 1998;144 (Pt 10):2905–14.

31. Taylor CM, Goldrick M, Lord L, Roberts IS. Mutations in the waaR gene of Escherichia coli which disrupt lipopolysaccharide outer core biosynthesis affect cell surface retention of group 2 capsular polysaccharides. J Bacteriol. 2006;188(3):1165–8.

32. Diao J, Bouwman C, Yan D, Kang J, Katakam AK, Liu P, et al. Peptidoglycan Association of Murein Lipoprotein Is Required for KpsD-Dependent Group 2 Capsular Polysaccharide Expression and Serum Resistance in a Uropathogenic. mBio. 2017;8(3).

33. Jiménez N, Senchenkova SN, Knirel YA, Pieretti G, Corsaro MM, Aquilini E, et al. Effects of lipopolysaccharide biosynthesis mutations on K1 polysaccharide association with the Escherichia coli cell surface. J Bacteriol. 2012;194(13):3356–67.

34. Singh S, Wilksch JJ, Dunstan RA, Mularski A, Wang N, Hocking D, et al. LPS O Antigen Plays a Key Role in Klebsiella pneumoniae Capsule Retention. Microbiol Spectr. 2022;10(4):e0151721.

35. Fresno S, Jiménez N, Izquierdo L, Merino S, Corsaro MM, De Castro C, et al. The ionic interaction of Klebsiella pneumoniae K2 capsule and core lipopolysaccharide. Microbiology (Reading). 2006;152(Pt 6):1807–18.

36. Datsenko KA, Wanner BL. One-step inactivation of chromosomal genes in Escherichia coli K-12 using PCR products. Proc Natl Acad Sci U S A. 2000;97(12):6640–5.

37. Goltermann L, Zhang M, Ebbensgaard AE, Fiodorovaite M, Yavari N, Løbner-Olesen A, et al. Effects of LPS Composition in. Front Microbiol. 2022;13:877377.

38. Grozdanov L, Zähringer U, Blum-Oehler G, Brade L, Henne A, Knirel YA, et al. A single nucleotide exchange in the wzy gene is responsible for the semirough O6 lipopolysaccharide phenotype and serum sensitivity of Escherichia coli strain Nissle 1917. J Bacteriol. 2002;184(21):5912–25.

39. Muheim C, Bakali A, Engström O, Wieslander Å, Daley DO, Widmalm G. Identification of a Fragment-Based Scaffold that Inhibits the Glycosyltransferase WaaG from Escherichia coli. Antibiotics (Basel). 2016;5(1).

40. Pagnout C, Sohm B, Razafitianamaharavo A, Caillet C, Offroy M, Leduc M, et al. Pleiotropic effects of rfa-gene mutations on Escherichia coli envelope properties. Sci Rep. 2019;9(1):9696.

41. Herlax V, de Alaniz MJ, Bakás L. Role of lipopolysaccharide on the structure and function of alpha-hemolysin from Escherichia coli. Chem Phys Lipids. 2005;135(2):107–15.

42. Phanphak S, Georgiades P, Li R, King J, Roberts IS, Waigh TA. Super-Resolution Fluorescence Microscopy Study of the Production of K1 Capsules by Escherichia coli: Evidence for the Differential Distribution of the Capsule at the Poles and the Equator of the Cell. Langmuir. 2019;35(16):5635–46.

43. Manzoni M, Rollini M, Piran E, Parini C. Preliminary characterisation of an Escherichia coli K5 lyase-deficient strain producing the K5 polysaccharide. Biotechnol Lett. 2004;26(4):351–6.

44. Maffei E, Shaidullina A, Burkolter M, Heyer Y, Estermann F, Druelle V, et al. Systematic exploration of Escherichia coli phage-host interactions with the BASEL phage collection. PLoS Biol. 2021;19(11):e3001424.

45. Gorzynski M, De Ville K, Week T, Jaramillo T, Danelishvili L. Understanding the Phage-Host Interaction Mechanism toward Improving the Efficacy of Current Antibiotics in. Biomedicines. 2023;11(5).

46. Chen Y, Li X, Wang S, Guan L, Hu D, Gao D, et al. A Novel Tail-Associated O91-Specific Polysaccharide Depolymerase from a Podophage Reveals Lytic Efficacy of Shiga Toxin-Producing Escherichia coli. Appl Environ Microbiol. 2020;86(9).

