## Supplementary file for "Synergy between Group 2 capsules and lipopolysaccharide underpins serum resistance in Extra-intestinal Pathogenic *Escherichia coli*"

| **Strain** | **Genotype/ Description** | **Selectable marker** | **Reference** |
| --- | --- | --- | --- |
| ***Escherichia coli*** |  |  |  |
| CFT073 Wild-type | Prototypic urosepsis isolate; O6:K2:H1 serotype | None | (1, 2) |
| CFT073 Δ*ksl* | *ksl::kan* | Kan^R^ | (3) |
| CFT073 Δ*ksl::FRT* | *ksl::FRT* | None | This study |
| CFT073 Δ*waaL* | *waaL::kan* | Kan^R^ | This study |
| CFT073 Δ*waaL::FRT* | *waaL::FRT* | None | This study |
| CFT073Δ*ksl ΔwaaL* | *ksl::gent, waaL::kan* | Gent^R^, Kan^R^ | This study |
| CFT073Δ*ksl::FRT ΔwaaL::FRT* | *ksl::FRT waaL::FRT* | None | This study |
| CFT073 Δ*wzy* | *wzy::kan* | Kan^R^ | This study |
| CFT073 Δ*wzy::FRT* | *wzy::FRT* | None | This study |
| CFT073 Δ*waaG* | *waaG::gent* | Gent^R^ | This study |
| CFT073 Δ*waaG::FRT* | *waaG::FRT* | None | This study |
| DH5α | K-12, serum sensitive control, *fhuA2 Δ(argF-lacZ)U169 phoA glnV44 Φ80 Δ(lacZ)M15 gyrA96 recA1 relA1 endA1 thi-1 hsdR17* | None | New England Biolabs |

**Table 2.1 Strains used in this study.** Kan - Kanamycin, Gent – Gentamicin.

| **Plasmids** | **Description** | **Resistance** | **Reference** |
| --- | --- | --- | --- |
| pCP20 | Possesses FLP flip recombinase gene, 30°C temperature-sensitive replication | Amp^R^ | (4) |
| pKD4 | Possesses FRT-flanked kanamycin resistance cassette | Kan^R^ | (5) |
| pMH2 | Possesses gentamicin resistance cassette which was utilised in mutant construction | Gent^R^ | Hunt *et al.*, 2015, Unpublished |
| pKD46 | Used to construct mutants through homologous recombination, possesses λ-Red recombinase genes *exo bet* and *gam* which are induced by L-arabinose, temperature sensitive replication at 30°C | Amp^R^ | (5) |
| pBWB536 | Complementation of O6-antigen synthesis genes | Amp^R^ | (6) |
| pXLW36 | pWKS30 with *kslCDABE* for complementation of *ksl* mutant | Amp^R^ | (7) |
| pBAD/His-WaaG | pBAD backbone; Vector used for complementation of *waaG* mutations. Under control of P*araBAD* arabinose-inducible promoter | Amp^R^ | (8) |

**Table 2.2 Plasmids used in this study. .** Kan - Kanamycin, Gent - Gentamicin, Amp – Ampicillin.

| Name | | | | Sequence (5’-3’) | | Purpose |
| --- | --- | --- | --- | --- | --- | --- |
|  | | 1 | | ATGTCGTTTTGTTGGAATGAAATTAACTCTGGTGTCAAGTCTTTAATTCTAT**TGTGTAGGCTGGAGCTGC** | **Amplify Kan^R^** cassette from pKD4 with homology to *waaL* | |
|  | | 2 | | TTACTTATCTAATAAACATTGGTCCGATTGTACTTTAAAATAAGCACAAAGGTC**CATATGAATATCCTCC** | **Amplify Kan^R^** cassette from pKD4 with homology to *waaL* | |
|  | | 3 | | GAGTCATTTGCGCACGAAAG | Screening primers for *waaL* in wild-type and mutant | |
|  | | 4 | | AGATGGTTTGTAGGGCTCCG | Screening primers for *waaL* in wild-type and mutant | |
|  | | 5 | | TAATGACGCAATTAAGTTATATCAAAATGATGAAAATGATGAAAATTTGAACATTTAGTATT**GTGTAGGCTGGAGCTGC** | Amplify Kan^R^ cassette from pKD4 with homology to *kpsT* | |
|  | | 6 | | GGGTATGAATAAAGATTTTTTGTTTGGATCAAAGTCAATATCATAATTAGGTC**CATATGAATATCCTCC** | Amplify Kan^R^ cassette from pKD4 with homology to *kpsS* | |
|  | | 7 | | GTCTTTATCAGAATATTAATGACGCAATTAAGTTATATCAAAATGATGAAAATTTGAACATTT**AGTGCGAATCCATGTGGGAGTTTA** | **Amplify Gentamicin^R^** cassette from pMH2 -homology to *kpsT* | |
|  | | 8 | | GAATGCATTGGGTATGAATAAAGATTTTTTGTTTGGATCAAAGTCAATATCATAATTTA**TTAGGTGGCGGTACTTGGGT** | **Amplify Gentamicin^R^** cassette from pMH2 -homology to *kpsS* | |
|  | | 9 | | CCCTGGTATGAAGCACGTTG | Screening primers for *ksl* operon in wild-type and mutant | |
|  | | 10 | | CATGTCGTGGAGTTAAGCCG | Screening primers for *ksl* operon in wild-type and mutant | |
|  | | 11 | | CGAATCCATGTGGGAGTTTA | Amplify Gentamicin^R^ cassette from pMH2 | |
|  | | 12 | | TTAGGTGGCGGTACTTGGGT | Amplify Gentamicin^R^ cassette from pMH2 | |
|  | | 13 | | TTGCCTTCCAGGCTGTTATC | *rplT* housekeeping control RT-PCR | |
|  | | 14 | | CTGCTTTCGCTTTTTCAACC | *rplT* housekeeping control RT-PCR | |
|  | | 15 | | CCCGTCATACTGACTGAGTACAT | *ksl2A* (region 2 capsule gene) RT-PCR | |
|  | | 16 | | TGCGGTGATTTGCAGTATCC | *ksl2A* (region 2 capsule gene) RT-PCR | |
|  | | 17 | | CCAGAGATTAACGCGCTCTAC | *waaQ* (R1 core biosynthesis) RT-PCR | |
|  | | 18 | | TGGCACGTAATACCTTGATGAG | *waaQ* (R1 core biosynthesis) RT-PCR | |
|  | | 19 | | GCTCAGCAATAGCCTCGCCGCAATTGGCGTCGACAATATA**CGAATCCATGTGGGAGTTTA** | *waaG:****gent*** mutagenesis F | |
|  | | 20 | | TCGATAAATTACTTCCCTCCTCCACGACAGGTACGTCGTT**TTAGGTGGCGGTACTTGGGT** | *waaG:****gent*** mutagenesis R | |
|  | | 21 | | GCAATGAAGATTGCGTTAAC | *waaG* screen F | |
|  | | 22 | | AGCGTGACCGAAATGAGATG | *waaG* screen R | |

**Table 2.3 Oligonucleotides used in this study.**

References

1.	Welch RA, Burland V, Plunkett G, Redford P, Roesch P, Rasko D, et al. Extensive mosaic structure revealed by the complete genome sequence of uropathogenic Escherichia coli. Proc Natl Acad Sci U S A. 2002;99(26):17020-4.

2.	Guyer DM, Kao JS, Mobley HL. Genomic analysis of a pathogenicity island in uropathogenic Escherichia coli CFT073: distribution of homologous sequences among isolates from patients with pyelonephritis, cystitis, and Catheter-associated bacteriuria and from fecal samples. Infect Immun. 1998;66(9):4411-7.

3.	Miajlovic H, Cooke NM, Moran GP, Rogers TR, Smith SG. Response of extraintestinal pathogenic Escherichia coli to human serum reveals a protective role for Rcs-regulated exopolysaccharide colanic acid. Infect Immun. 2014;82(1):298-305.

4.	Cherepanov PP, Wackernagel W. Gene disruption in Escherichia coli: TcR and KmR cassettes with the option of Flp-catalyzed excision of the antibiotic-resistance determinant. Gene. 1995;158(1):9-14.

5.	Datsenko KA, Wanner BL. One-step inactivation of chromosomal genes in Escherichia coli K-12 using PCR products. Proc Natl Acad Sci U S A. 2000;97(12):6640-5.

6.	Sarkar S, Ulett GC, Totsika M, Phan MD, Schembri MA. Role of capsule and O antigen in the virulence of uropathogenic Escherichia coli. PLoS One. 2014;9(4):e94786.

7.	Buckles EL, Wang X, Lane MC, Lockatell CV, Johnson DE, Rasko DA, et al. Role of the K2 capsule in Escherichia coli urinary tract infection and serum resistance. J Infect Dis. 2009;199(11):1689-97.

8.	Muheim C, Bakali A, Engström O, Wieslander Å, Daley DO, Widmalm G. Identification of a Fragment-Based Scaffold that Inhibits the Glycosyltransferase WaaG from Escherichia coli. Antibiotics (Basel). 2016;5(1).
